# The inhibition of LSD1 via sequestration contributes to tau-mediated neurodegeneration

**DOI:** 10.1101/745133

**Authors:** Amanda K. Engstrom, Alicia C. Walker, Rohitha A. Moudgal, Dexter A. Myrick, Stephanie M. Kyle, Yu Bai, M Jordan Rowley, David J. Katz

## Abstract

Tauopathies are a class of neurodegenerative diseases associated with pathological tau. Despite many advances in our understanding of these diseases, the direct mechanism through which tau contributes to neurodegeneration remains poorly understood. Previously, our lab implicated the histone demethylase LSD1 in tau-induced neurodegeneration by showing that LSD1 localizes to pathological tau aggregates in Alzheimer’s disease cases, and that it is continuously required for the survival of hippocampal and cortical neurons in mice. Here, we utilize the P301S tauopathy mouse model to demonstrate that pathological tau can exclude LSD1 from the nucleus in neurons. In addition, we show that reducing LSD1 in these mice is sufficient to highly exacerbate tau-mediated neurodegeneration and tau-induced gene expression changes. Finally, we find that overexpressing LSD1 in the hippocampus of tauopathy mice, even after pathology has formed, is sufficient to significantly delay neurodegeneration and counteract tau-induced expression changes. These results suggest that inhibiting LSD1 via sequestration contributes to tau-mediated neurodegeneration. Thus, LSD1 is a promising therapeutic target for tauopathies such as Alzheimer’s disease.

**SIGNIFICANCE STATEMENT:** We have made the novel discovery that pathological tau functions through the histone demethylase LSD1 in the Alzheimer’s disease pathway. Thus, we have identified a mechanism that links tau to the downstream neuronal dysfunction pathways. This step can potentially be targeted therapeutically, after the onset of dementia symptoms, to block the progression of dementia in Alzheimer’s disease patients.

## INTRODUCTION

Tauopathies such as corticobasal degeneration, progressive supranuclear palsy, and frontotemporal lobar degeneration with tau inclusions are neurodegenerative diseases pathologically defined by different forms of tau positive intraneuronal deposits (*1–5*). In addition to these primary tauopathies, neuropathological observations of postmortem Alzheimer’s disease (AD) brains show the presence neurofibrillary tangles (NFTs) of hyperphosphorylated tau protein, as well as plaques containing β-amyloid (Aβ) peptide (*6–9*). AD is the leading cause of age-related dementia, resulting from neuronal cell death in the frontal and temporal cortices, as well as the hippocampus (*7*). As dementia progresses, the spatial pattern of tau pathology highly correlates with the level of cognitive impairment (*10–13*). In addition, Aβ oligomers and/or plaques can enhance tau pathology in various mouse models (*14, 15*), and there is increasing evidence that accumulation of Aβ plaques can contribute to tau pathology (*3, 16, 17*). The most well-defined physiological role of tau is in stabilizing microtubules, particularly in neuronal axons (*2*). However, in the pathological state, tau becomes aberrantly phosphorylated (*2, 18, 19*), truncated (*1, 4*), and aggregates into oligomers and larger insoluble filaments (*20, 21*). This pathology is thought to trigger synaptic loss, dramatic genome-wide expression changes, increased inflammatory response, and neuronal cell death (*22–25*). These data suggest that pathological tau may be a downstream mediator of the neurotoxic effects leading to neuronal degeneration in AD.

Previously, our lab demonstrated that deleting the histone demethylase *Lsd1* in adult mice leads to significant neuronal cell death in the hippocampus and cortex with associated learning and memory defects (*26*). In this mouse model, loss of *Lsd1* induces genome-wide expression changes that significantly overlap with those observed in the brains of postmortem human AD cases, but not other neurodegenerative diseases, such as Parkinson’s disease or amyotrophic lateral sclerosis (ALS) cases. Consistent with this overlap, we observed LSD1 protein mislocalized to cytoplasmic NFTs, but not associated with Aβ plaques in AD cases or Lewy bodies of α-synuclein in Parkinson’s disease cases (*26*). In control cases, LSD1 remains strictly confined to the nucleus (*26*), due to it’s well-defined nuclear localization signal (*27*). These data highlight the requirement for LSD1 in neuronal survival and suggest that the nuclear function of the histone demethylase LSD1 could be disrupted by mislocalization to pathological tau.

To investigate how LSD1 may contribute to tau-mediated neurodegeneration, we utilized the PS19 P301S tauopathy mouse model (hereafter referred to as PS19 Tau). PS19 Tau mice express a P301S mutated form of the human tau protein, originally identified in a frontotemporal dementia with parkinsonism (FTDP-17) patient, driven by the prion promoter throughout the nervous system (*28*). When expressed in mice, the P301S tau protein is prone to hyperphosphorylation and somatodendritic aggregation, without the presence of Aβ plaques. PS19 Tau mice develop a heavy pathological tau burden and have been well characterized for the temporal progression of tau pathology and disease-related phenotypes (*29, 30*). However, the mechanism of neuronal cell death caused by pathological tau is still unknown.

Here, we provide functional data that the inhibition of LSD1 function contributes to tau induced neurodegeneration. We demonstrate in PS19 Tau mice that pathological tau sequesters LSD1 in the cytoplasm of neurons throughout the brain. This results in depletion of LSD1 from the nucleus. Additionally, we provide genetic and molecular evidence that pathological tau contributes to neurodegeneration by disrupting LSD1. Finally, we show that overexpressing LSD1 in hippocampal neurons is sufficient to suppress neuronal cell death even after pathological tau has formed. We propose that pathological tau contributes to neuronal cell death by sequestering LSD1 in the cytoplasm, depleting the nuclear pool of LSD1 that is required for neuronal survival.

## RESULTS

### Tau pathology depletes LSD1 from the nucleus in the PS19 Tau mouse model

Previously, we showed in human AD cases that LSD1 protein inappropriately colocalizes with NFTs in the cell body of hippocampal and cortical neurons, while in unaffected controls LSD1 was properly localized exclusively to the nucleus (*26*). However, because neurons in AD cases with intracellular NFTs presumably die and are cleared, it was difficult to determine whether tau prevents LSD1 from localizing to the nucleus in a dying neuron. To address this possibility, we performed LSD1 immunofluorescence on 12 month old PS19 Tau mice, which have significant tau pathology (*28*). Because PS19 Tau mice undergo neurodegeneration over a shortened period, there are more neurons undergoing neurodegeneration at any given time point. Thus, we reasoned that it may be possible to observe LSD1 depletion from the nucleus. Similar to what we observe in humans, LSD1 protein in 12 month old Wild Type mice was strictly localized to the nucleus of neurons in the cerebral cortex (Fig. 1A-C) and the hippocampus (Fig. S1 A-C). This was expected since LSD1 has a well-defined nuclear localization signal (*27*). However, in 12 month old PS19 Tau mice, LSD1 protein was sequestered in the cytoplasm and depleted from the nucleus both in the cerebral cortex (Fig. 1D-F,M,N) and the hippocampus (Fig. S1 D-F). These are both regions where we observe substantial cytoplasmic tau pathology (Fig. 1G-I; Fig. S1 G-I). Similarly, in other brain regions that accumulate tau pathology, such as the thalamus and amygdala, LSD1 was localized to the nucleus in 12 month old Wild Type control mice (Fig. S1 J-O), but abnormally localized to the cytoplasm in PS19 Tau mouse littermates (Fig. S1 P-U). Overall, we observed sequestration of LSD1 in 6 out of 7 mice analyzed. In each of the 6 mice, there were varying levels of sequestration ranging from LSD1 found in both the nucleus and cytoplasm (Fig. S1 V-X, M,N), to depletion from the nucleus (Fig. S1 Y-DD, M,N). The mouse that has little sequestration (denoted by the ^ in Fig. 1M) has extensive tau pathology, but this pathology is largely extracellular rather than cytoplasmic, indicating that neurons with extensive LSD1 sequestration may have died and been cleared.

**Fig. 1:**
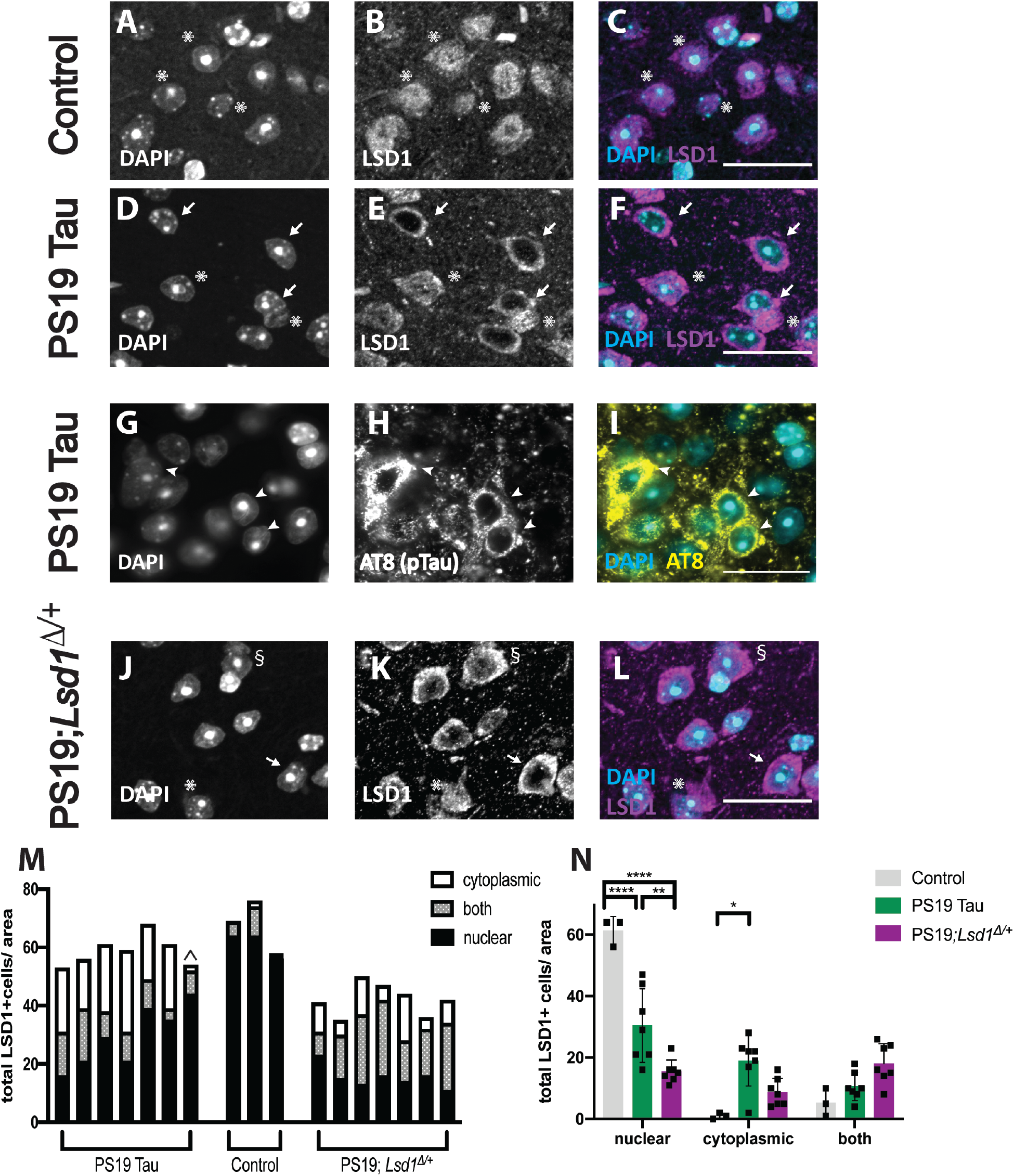
LSD1 sequestration and tau accumulation in the presence of pathological tau. **A-C**, Representative immunofluorescence of 12 month old control Wild Type mice showing DAPI (**A**), LSD1 (**B**), and merged (**C**) in the cerebral cortex where LSD1 is localized specifically to DAPI positive nuclei. **D-F**, Representative image of the cerebral cortex in 12 month old PS19 Tau mice. Staining for DAPI (**D**), LSD1 (**E**), and merged (**F**) shows that LSD1 is localized outside the nucleus, and depleted from the DAPI positive nucleus. Arrows denote cells where LSD1 is localized outside of the nucleus, and asterisks denote LSD1 localized specifically to the nucleus. **G-I,** Representative immunofluorescence of 12 month old PS19 Tau mouse with staining for DAPI (**G**), AT8 positive hyperphosphorylated tau (**H**) and merge (**I**) where hyper-phosphorylated tau accumulates in the cytoplasm of the cell body. Arrowheads denote hyper-phosphorylated tau. **J-L,** Representative immunofluorescence of 12 month old control Tau mice that are heterozygous for *Lsd1* (PS19; *Lsd1*^*Δ*/+^) showing DAPI (**J**), LSD1 (**K**), and merged (**L**) in the cerebral cortex. Arrows denote cells where LSD1 is localized outside of the nucleus, asterisks denote LSD1 localized specifically inside the nucleus, and § denotes cells where LSD1 is both nuclear and cytoplasmic. Scale bars=25μm. **M,N**, Quantification of the LSD1 localization represented in **A-F, J-L**.(**M**) Each bar represents a single mouse. LSD1 was scored as being localized to the nucleus, the cytoplasm, or both areas. ^ denotes the one PS19 Tau mouse with low levels of observed cytoplasmic LSD1 localization. (**N**) Combined analysis from **M** in each cell region for Control (WT *n*=3), PS19 Tau (*n*=7), and PS19;*Lsd1*^*Δ*/+^ (*n*=7) mice. Values are mean ± SEM, two-way analysis of variance (ANOVA) with Tukey’s post hoc test *P<0.05, **P<0.01,****P<0.001.

### Reduction of LSD1 increases the mouse tauopathy phenotype

If the presence of pathological tau in the cytoplasm is leading to neuronal cell death through the sequestration and nuclear depletion of LSD1, we would expect that lowering the overall levels of LSD1 would accelerate depletion and exacerbate the progression of disease. To test this, we made PS19 Tau mice heterozygous for *Lsd1* (hereafter referred to as PS19;*Lsd1*^*Δ*/+^, Fig. S2 A). *Lsd1* was deleted using *Vasa-Cre* (*31*). The *Lsd1* deletion created when the floxed allele recombines is a null allele that has previously been characterized (*26, 32, 33*). Once the *Lsd1* deletion passes through the germline, *Lsd1* is heterozygous throughout the mouse. These *Lsd1* heterozygotes (hereafter referred to as *Lsd1*^*Δ*/+^) have a functioning copy of *Lsd1* and don’t have any phenotypes associated with the heterozygous loss of LSD1 function (*33–35*). Thus, *Lsd1* heterozygosity does not completely compromise LSD1 function. Instead mice that are heterozygous for *Lsd1* are sensitized to mechanisms affecting LSD1 localization and function. Consistent with this, we observed a 20% reduction in transcript levels (Fig. S2 B) and a 35% reduction in protein levels (Fig. S2 C,D) in *Lsd1*^*Δ*/+^ mice compared to their *Lsd1*^+/+^ littermates. Surprisingly, PS19 Tau mice have a 20% increase in LSD1 protein levels compared to *Lsd1*^+/+^ littermates. It is possible that this is due to some type of compensation because LSD1 is being sequestered. Nevertheless, consistent with the reduction in LSD1 that we observe in *Lsd1*^*Δ*/+^ mice, PS19;*Lsd1*^*Δ*/+^ mice have a similar 14% reduction in transcript levels (Fig. S2 B) and a 31% reduction in protein levels (Fig. S2 C,D) compared to PS19 Tau littermates. This enables us to directly assess the effect of reducing LSD1 on tau-induced neurodegeneration. In addition, all genotypes were born at normal Mendelian ratios with equal male/female ratios.

To determine if reducing LSD1 protein levels accelerates tau-induced depletion of LSD1 from the nucleus, we analyzed localization of LSD1 in PS19;*Lsd1*^*Δ*/+^ mice versus PS19 Tau controls. When one copy of *Lsd1* was removed from PS19 Tau mice, there was a significant decrease in the nuclear localization of LSD1 compared to PS19 Tau mice (Fig. 1J-N), accompanied by an increase in the number of cells that have LSD1 in both the nucleus and the cytoplasm (Fig 1J-N). This suggests that reducing LSD1 enables tau to accelerate the depletion of LSD1.

Since lowering LSD1 accelerates tau-induced depletion of LSD1 from the nucleus, we determined if this acceleration exacerbates the progression of disease, by analyzing survival, paralysis, neurodegeneration and genome-wide gene expression changes in PS19;*Lsd1*^*Δ*/+^ mice versus PS19 Tau controls. As expected, *Lsd1*^*Δ*/+^ mice had normal survival (Fig. 2A). In contrast, PS19 Tau mice had a reduced overall survival (Fig. 2A) (*28*). When one copy of *Lsd1* was removed from PS19 Tau mice, their reduced survival was significantly exacerbated (P-value = 0.0017, Fig. 2A). As expected, there was little effect on the onset of reduced viability. The initial decline in the survival of PS19;*Lsd1*^*Δ*/+^ mice started only slightly earlier than PS19 Tau mice, but after the appearance of hyperphosphorylated (AT8 positive) pathological tau in neurons, as quantified by the average AT8 positive tau immunoreactivity per area in the hippocampus and cortex (Fig. S6 K-M)(*28*). This suggests that pathological tau may have to be present in order for *Lsd1* heterozygosity to have deleterious effects. Subsequently, PS19;*Lsd1*^*Δ*/+^ mice had a 14% reduction in median lifespan compared to PS19 Tau mice and reached median survival 44 days earlier then PS19 Tau mice. In addition, there was a further exacerbation of reduced lifespan as pathology became more severe. PS19;*Lsd1*^*Δ*/+^ mice reached the point when there was only 10% of the population remaining 83 days earlier than PS19 Tau mice, and all but one of the last 25% of PS19;*Lsd1*^*Δ*/+^ mice died between 11.5-13.5 months, compared to 13.5-19 months in PS19 Tau mice. As a result, 28% of PS19 Tau mice were still alive after all but one of the PS19;*Lsd1*^*Δ*/+^ mice had died (Fig. 2A).

**Fig. 2:**
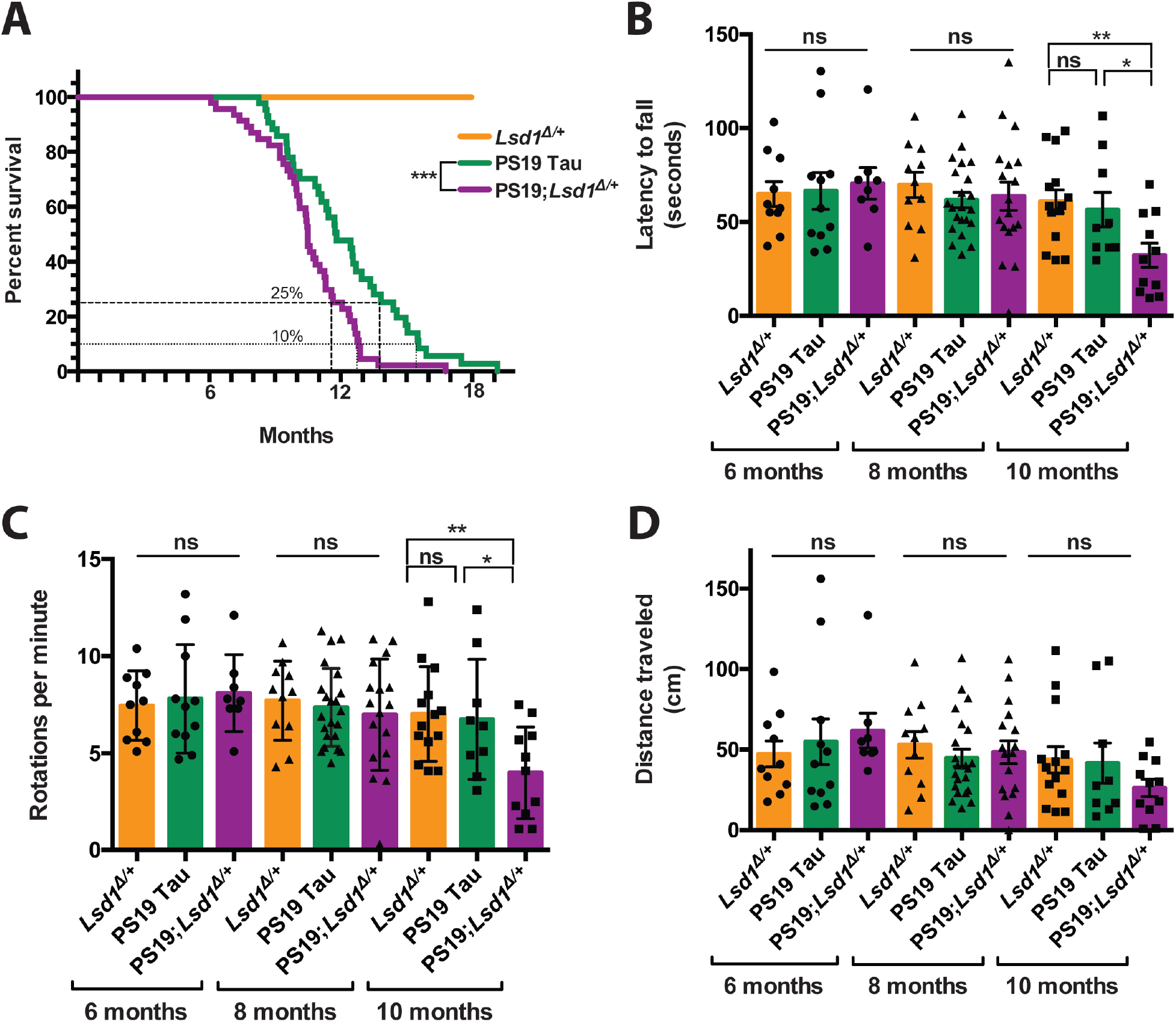
Reduction of *Lsd1* exacerbates the PS19 Tau mouse paralysis phenotype. **A,** Lifespan curve showing that no *Lsd1*^*Δ*/+^ mice died before 18 months (orange, *n*=20). PS19;*Lsd1*^*Δ*/+^ mice (purple, *n*=44) have a significant reduction in survival compared to PS19 Tau mice with wild-type levels of *Lsd1* (green, *n*=37)(Log-rank Mantle-Cox test ***P<0.005). **B-D**, Rotarod testing of latency to fall (in seconds) (**B**), rotations per minute (when the mouse fell) (**C**), and distance traveled (in centimeters) (**D**) for mice at age 6, 8 and 10 months. *Lsd1*^*Δ*/+^ (orange, *n*=10,11,14), PS19 Tau (green, *n*=11,22,9), and PS19;*Lsd1*^*Δ*/+^ (purple, *n*=8,17,11). Values are mean ± SEM (two-way analysis of variance (ANOVA) with Tukey’s post hoc test *P<0.05 **P<0.01, ns=not significant).

PS19 Tau mice develop paralysis starting with hind limb clasping which progresses until they are unable to feed (*28*). In our hands, PS19 Tau mice displayed intermittent hind limb clasping starting at approximately 6 months of age. At 12 months, these mice had a severe clasp, but were still mobile. This is delayed compared to what was originally reported by Yoshiyama and colleagues (*28*). PS19;*Lsd1*^*Δ*/+^ mice also displayed intermittent hind limb clasping beginning at approximately 6 months of age. However PS19;*Lsd1*^*Δ*/+^ mice became terminally paralyzed at a faster rate compared to PS19 Tau mouse littermates. At 12 months, when PS19 Tau mice were still mobile, PS19;*Lsd1*^*Δ*/+^ mice were severely paralyzed and typically terminal (Movie S1). To quantitatively assess paralysis we performed rotarod and grid performance tests. In the rotarod, we assessed the ability of the mice to stay on the rotating rod (latency to fall) (Fig. 2B), the speed of the rod at which they fall off the rotarod (rotations per minute) (Fig. 2C), and the total distance traveled (Fig. 2D). All genotypes performed the same at 6 months and 8 months (Fig. 2B-D). However, at 10 months, when PS19 Tau mice still performed normally, PS19;*Lsd1*^*Δ*/+^ mice had a significant deficit in mobility (P-value < 0.01, Fig. 2B,C). A deficit in PS19;*Lsd1*^*Δ*/+^ mice was also observed at 10 months in the total distance traveled (Fig. 2D) and in grid performance testing (Fig. S3 A), though neither were statistically significant.

To further investigate the exacerbation of paralysis we examined spinal cord motor neurons. Healthy motor neurons from *Lsd1*^+/+^ control mice express LSD1 (Fig. S3 B-D) and are classically identified by circular nuclei at the center of a large cell body. In contrast to the healthy motor neurons we observed in 12 month old PS19 Tau mice (Fig. S3 E), many of the motor neurons in PS19;*Lsd1*^*Δ*/+^ mice at 12 months had abnormal morphology, with the nucleus skewed to the edge of the cell body (Fig. S3 E vs. S3 F) and a ballooned cell body (Fig. S3 F). Within the cell body we found aberrant hyperphosphorylated NFH (heavy chain neurofilament), which is a sign of activated neuronal stress pathways (Fig S3 G vs. S3 H) (*36, 37*). This abnormal morphology is highly reminiscent of a well-established process known as chromatolysis, which is characterized by swelling of the neuronal cell body, disruption of Nissl granules, and pyknotic or shrunken nuclei abnormally skewed to the edge of the cell body (*38, 39*). Chromatolysis, which is linked to neuronal stress and often leads to apoptosis (*39*), has been observed in AD and other neurodegenerative diseases (*38, 40–42*).

### Reduction of LSD1 exacerbates PS19 Tau neurodegeneration

In addition to accelerating the paralysis phenotype, reducing the level of *Lsd1* in PS19 Tau mice exacerbated neuronal cell death in the brain. At 6 months and 8 months, we observed no difference between genotypes in the overall morphology in the hippocampus (Fig. S4 A-H), based on histological analysis. There was also no difference between *Lsd1*^+/+^ and *Lsd1*^*Δ*/+^ mice at 10 months or 12 months (Fig. 3A,B; Fig. S4 I,J,M,N; Fig. S5 A,B,E,F,I,J,M,N). At 10 months, PS19 Tau mice had very little cell loss in the hippocampus compared to *Lsd1*^+/+^ and *Lsd1*^*Δ*/+^ control mice (Fig. S4 I-K, M-O). In contrast, at 10 months, PS19;*Lsd1*^*Δ*/+^ mice had dramatic cell loss both in the CA1 region of the hippocampus and throughout the posterior hippocampus (Fig. S4 L,P). At 12 months, the PS19 Tau mice had a slight decrease in CA1 and CA3 neurons spanning the hippocampal pyramidal cell layer compared to *Lsd1*^+/+^ control mice (17% and 19.4% respectively, Fig. 3A,B; Fig. S5 A-C, E-G). In comparison, PS19;*Lsd1*^*Δ*/+^ mice had a 52% and 54% decrease in the CA1 and CA3 respectively (Fig. 3A,B; Fig. S5 D,H) compared to *Lsd1*^+/+^ control mice. This resulted in decreased overall brain size (Fig. 3C) and brain weight (Fig. 3D) in PS19;*Lsd1*^*Δ*/+^ mice compared to PS19 Tau and *Lsd1*^*Δ*/+^ mouse littermates. Additionally, at 12 months there were increased levels of cell loss in the Dentate Gyrus (Fig. S5 I-L), and throughout the posterior hippocampus (Fig. S5 M-P).

**Fig. 3:**
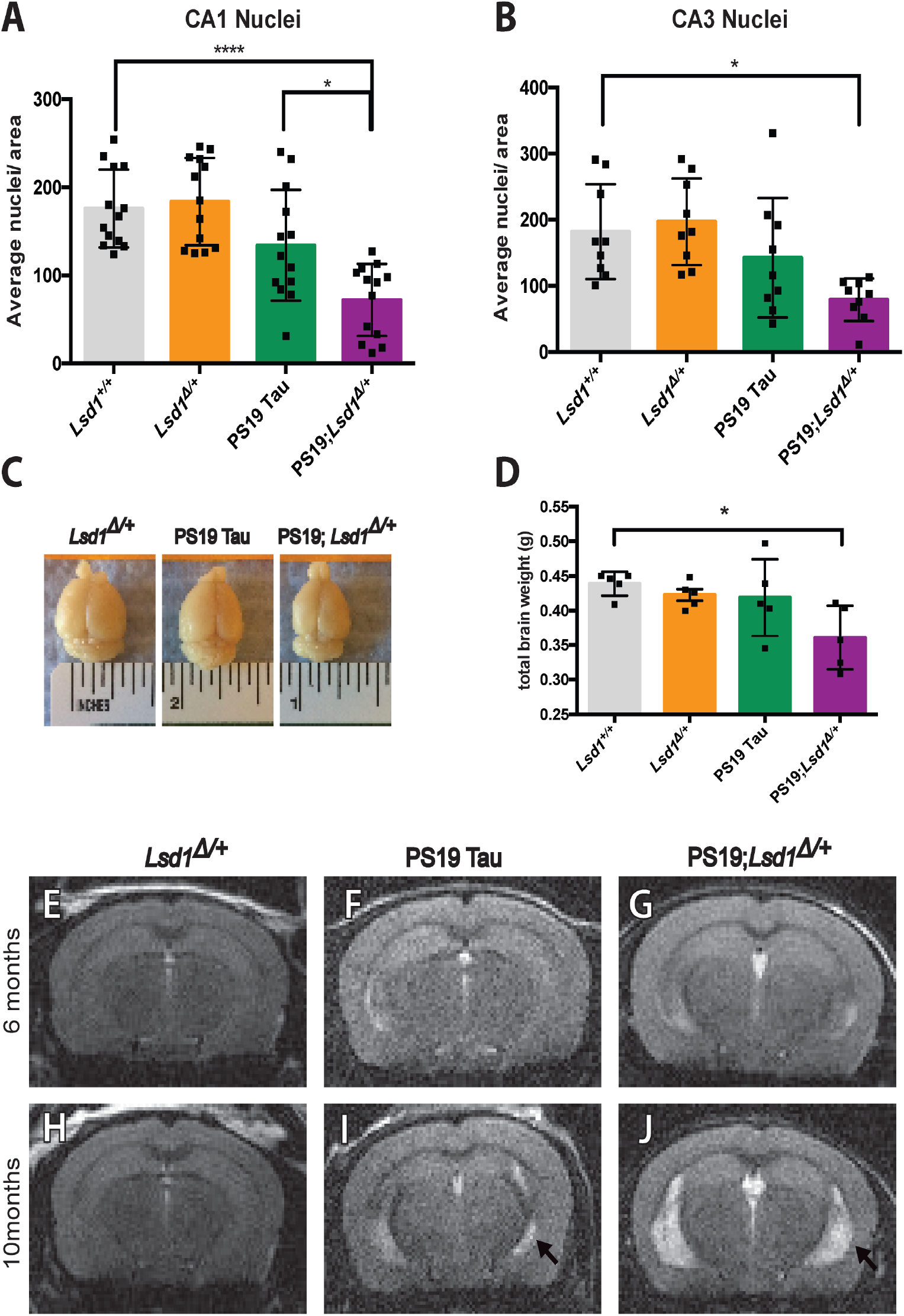
Reduction of *Lsd1* exacerbates neurodegeneration in PS19 Tau mice. **A,B,** Average nuclei per area in the CA1 (**A**) and CA3 (**B**) regions of the hippocampus in 12 month old *Lsd1*^+/+^, *Lsd1*^*Δ*/+^, PS19 Tau, and PS19;*Lsd1*^*Δ*/+^ mice. Quantification from histology represented in **Fig. S5 A-H**. Values are mean ± SD (**A**, *n*=13 & **B,** *n*=9). **C,** Representative image of the brains of 12 month old *Lsd1*^*Δ*/+^, PS19 Tau, and PS19;*Lsd1*^*Δ*/+^ littermates. **D,** Total brain weight of 12 month old littermates represented in **Fig. 3C.** Values are mean ± SD (*n*=5). For all graphs: one-way analysis of variance (ANOVA) with Tukey’s post hoc test (two-sided) *P<0.05,****P<0.001. **E-J,** Representative image of T2-weighted RARE coronal MRI taken from 6 months (**E-G**) and 10 months (**H-jJ**) of age in *Lsd1*^*Δ*/+^ (**E,H**), PS19 (**F,J**), and PS19;*Lsd1*^*Δ*/+^ (**G,J**) mice (*n*=3). Arrow denotes region of hippocampal atrophy.

Along with the histology, we monitored the progression of neuronal cell death in the same individual over time by performing magnetic resonance imaging (MRI) at 6 months and again at 10 months (Movie S2). At 6 months, there was no sign of cell loss or ventricular dilatation in *Lsd1*^*Δ*/+^, PS19 Tau, or PS19;*Lsd1*^*Δ*/+^ mice (Fig. 3E-G). However, at 10 months the MRI showed that there was dramatic ventricular dilatation in PS19;*Lsd1*^*Δ*/+^ mice, as evidenced by high-intensity areas in T2-weighted imaging, with substantial hippocampal and neocortical atrophy (Fig. 3J vs. 3H, Movie S2). At this timepoint, PS19 Tau mice had some ventricular dilatation and hippocampal atrophy throughout the hippocampus (Fig. 3I), but much less than PS19;*Lsd1*^*Δ*/+^ mice (Fig. 3J, Movie S2. 0:51sec vs. 1:00sec).

### Tau pathology is not affected by change in LSD1 levels

Since LSD1 is a chromatin regulator, it is possible that reducing LSD1 protein levels affects the PS19 Tau transgene. However, we confirmed that there was no difference between PS19 Tau mice and PS19;*Lsd1*^*Δ*/+^ mice in the endogenous mouse *Mapt* RNA expression, nor in the human P301S MAPT transgene expression (Fig. S6 A). It is also possible that LSD1 affects tau pathology. To test this, we performed immunohistochemistry staining for a hyperphosphorylated form of tau (AT8). As expected, we did not observe any AT8 positive staining in *Lsd1*^*Δ*/+^ at 6, 8, or 10 months (Fig. S6 B,E,H). Additionally, we observed very little AT8 positive immunoreactivity at 6 months in both PS19 Tau or PS19;*Lsd1*^*Δ*/+^ mice (Fig. S6 C,D). At 8 months both PS19 Tau and PS19;*Lsd1*^*Δ*/+^ mice had low but consistent levels of AT8 positive immunoreactivity (Fig. S6 F,G), and by 10 months both PS19 Tau and PS19;*Lsd1*^*Δ*/+^ mice developed the same high level of AT8 positive tau immunoreactivity (Fig. S6 I,J). This was consistent throughout both the CA1 and CA3 regions of the hippocampus and the cerebral cortex (Fig. S6 K-M). We also did not observe any difference between PS19 Tau mice and PS19;*Lsd1*^*Δ*/+^ mice when assaying PHF1 (an alternative phospho-tau antibody) immunoreactivity in the CA1 region of the hippocampus at 8 months (Fig. S7 A-C, G) and 10 months (Fig. S7 D-F, G), nor in the CA3 region of the hippocampus or the cerebral cortex at 8 and 10 months (Fig. S7 H,I).

### The functional interaction between tau pathology and LSD1 inhibition is specific

To test the specificity of the functional interaction between tau pathology and LSD1, we investigated the overlap in the effected molecular pathways associated with both pathological tau and *Lsd1* heterozygosity. To address this, we performed RNA sequencing on the hippocampus of 9 month *Lsd1*^+/+^, *Lsd1*^*Δ*/+^, PS19 Tau, and PS19;*Lsd1*^*Δ*/+^ littermates. As opposed to analyzing transcriptional changes at the terminal stage of disease, this time point allows us to assess molecular changes prior to the onset of neuronal cell death. This is also the time point that we observed the earliest signs of exacerbation of paralysis in PS19;*Lsd1*^*Δ*/+^ mice. Because of this early stage in the progression of the disease, we would not expect dramatic changes in transcription overall. Nevertheless, if tau pathology is inhibiting LSD1 function, we would expect that the genome-wide expression changes induced by tau might be exacerbated by a reduction in LSD1. The RNA-seq analysis detected 112 significant gene expression changes in PS19 Tau mice compared to *Lsd1*^+/+^ (Fig S8 A,B), and 295 significant gene expression changes in PS19;*Lsd1*^*Δ*/+^ mice compared to *Lsd1*^+/+^ (Fig S8 C,D). Importantly, *Lsd1*^*Δ*/+^ mice had only 4 gene expression changes observed genome-wide (Fig. S8 E,F), indicating that the partially reduced level of LSD1 expression had very little effect on transcription on its own. This is consistent with the lack of phenotype in these animals.

We first examined the relationship between tau-induced expression changes and the effects of *Lsd1* heterozygosity by comparing the transcriptional changes observed in PS19 Tau mice with PS19;*Lsd1*^*Δ*/+^ mice. In PS19 Tau mice that do not yet have significant neurodegeneration, we identified 112 genes (107 up and 5 down) that were differentially expressed. All of these 112 genes were changed in the same direction in PS19;*Lsd1*^*Δ*/+^ mice. In addition, amongst the 112 genes changed in the same direction, 104 (93%) had exacerbated expression in PS19;*Lsd1*^*Δ*/+^ mice compared to PS19 Tau mice (Fig. 4A). Based on this overlap, we further compared the expression changes between PS19 Tau mice and PS19;*Lsd1*^*Δ*/+^ mice genome-wide. Amongst the transcripts that were changed in both PS19 Tau mice and PS19;*Lsd1*^*Δ*/+^ mice compared to *Lsd1*^+/+^ mice, 77% changed in the same direction (either up or down). Consistent with this overlap in gene expression, Gene Set Enrichment Analysis demonstrated that the pathways that are affected in both PS19 Tau mice and PS19;*Lsd1*^*Δ*/+^ mice are very similar (Fig. S8 G-J). However, the genes affected in both sets of mice tended to be further exacerbated in PS19;*Lsd1*^*Δ*/+^ mice compared to PS19 Tau mice. Amongst the 77% of genes that changed in the same direction in both PS19 Tau mice and PS19;*Lsd1*^*Δ*/+^ mice, 75% of these transcripts had a higher fold-change in PS19;*Lsd1*^*Δ*/+^ mice compared to PS19 Tau mice (Fig. 4B). Importantly, the gene expression pathways that are induced by the PS19 transgene overlap substantially with those observed in human AD case (*43*). This suggests that reduction of *Lsd1* exacerbates human AD pathways.

**Fig. 4:**
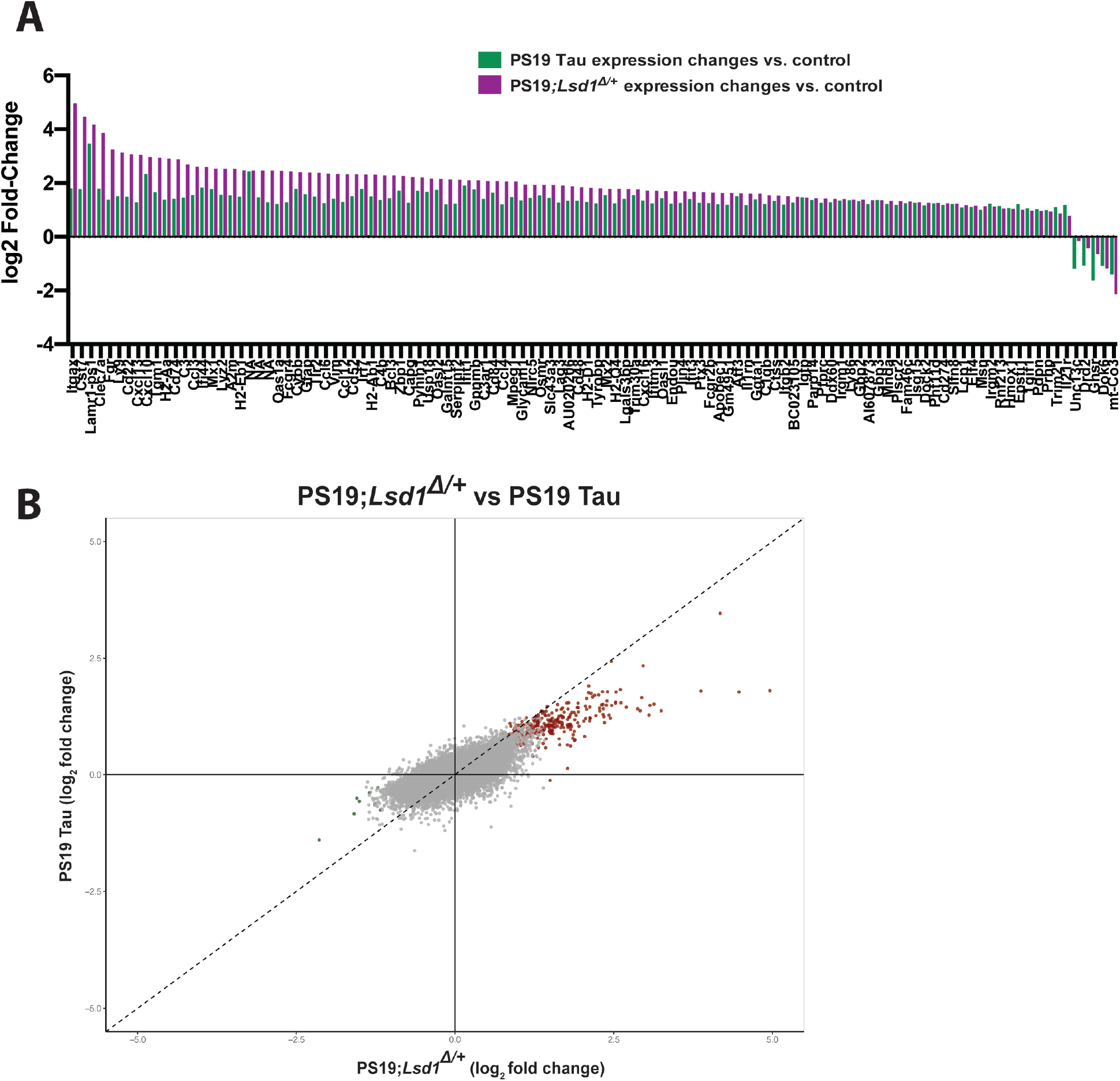
Molecular overlap between loss of LSD1 function and tauopathy. **A,** Histogram (log_2_ fold change) of the 112 genes that have significant changes in gene expression in the PS19 Tau mice (green, *n*=4) and their corresponding gene expression change in PS19;*Lsd1*^*Δ*/+^ mice (purple, *n*=*4*). **B,** Scatter plot showing the correlation between the genome-wide log_2_ fold change in gene expression between PS19 Tau and PS19;*Lsd1*^*Δ*/+^. The most significantly changed genes in PS19;*Lsd1*^*Δ*/+^ mice are shown in red (upregulated) and green (downregulated). All other genes are shown in grey. Dotted line represents 1:1 relationship between gene expression changes in PS19 Tau vs. PS19;*Lsd1*^*Δ*/+^. Exacerbated genes fall to the right of the dotted line in the positively correlated quadrant and to the left of the dotted line in the negatively correlated quadrant. Genes with correlated expression changes are found in the top right and bottom left quadrants, while genes that do not correlate are found in the opposite quadrants.

### Overexpression of LSD1 rescues neurodegeneration in the hippocampus of PS19 Tau mice

Our data demonstrate that reduction of LSD1 protein exacerbates the tauopathy phenotype in PS19 Tau mice. Based on this, we considered the possibility that overexpression of LSD1 might counter the loss of LSD1 from the nucleus and protect against neurodegeneration in PS19 Tau mice. To address this possibility, we injected PS19 Tau mice with a neuronal specific virus (AAV-DJ driven by the synapsin promoter) expressing either the full length LSD1 protein with an N-terminal HA tag (hereafter referred to as PS19-LSD1 inj) or a control virus expressing only the HA tag (hereafter referred to as PS19-HA inj). Additionally, to control for the effects of viral injection, we injected Wild Type littermates with the HA only expressing virus (hereafter referred to as WT-HA inj). All injections were performed directly into the hippocampus at 8-8.5 months, when tau pathology is already present throughout the nervous system. Immunolabeling for the HA tag demonstrated that the virus is specific to NeuN+ neurons (Fig. S9 A-D), with no HA expression observed in IBA+ microglia (Fig. S9 E-H), or GFAP+ astrocytes (Fig. S9 I-L). It also confirmed that virally expressed LSD1 is nuclear (Fig. S9 M) and confined to the hippocampus (Fig. S9 N), even at 11-11.5 months, when LSD1 is normally becoming sequestered to the cytoplasm (Fig. S9 O-R). After 3 months of overexpression, 11-11.5 month old mice were euthanized, and the brains were analyzed. Injections resulted in a ~6-fold increase in expression of LSD1 in the hippocampus compared to endogenous LSD1 in the PS19-HA inj mice, but no increase in the cerebellum (Fig. S9 S,T). As expected, because the viral injections were restricted to the hippocampus, the mice injected with LSD1 still developed paralysis (Movie S3). This confirms that the tau transgene is expressed and functioning. Additionally, we did not observe a difference in total levels of AT8 positive tau immunoreactivity (Fig. S9 U-X). Therefore, any modulation to the phenotype was not due to changes in tau pathology.

Injected mice were evaluated for cell death by neuronal cell counts in the hippocampus. Injection of LSD1 virus into the hippocampus of PS19 Tau mice rescued the neurodegeneration phenotype. At 11 months, compared to WT-HA inj control mice, 70% of the PS19-HA inj mice had hippocampal cell counts that were below the lowest WT-HA inj control, while none of the Tau mice injected with the LSD1 virus were below this level. Overall, we observed significantly more neurons (P-value <0.05) spanning the pyramidal cell layer (84% of WT-HA inj CA1 counts) compared to their PS19-HA inj littermates (59% of WT-HA inj CA1 counts), such that overall the neuronal cell count in PS19-LSD1 inj mice was not statistically different from the WT-HA inj (Fig. 5A-D). Additionally, in the histological analysis we observed a large number of cells infiltrating the hippocampus in PS19-HA inj mice compared to WT-HA inj littermates (Fig. 5B vs. 5C). Marker analysis demonstrated that this was due to a strong inflammatory response, with a large increase in the number of GFAP+ astrocytes (Fig. S10 A-C vs. D-F) and TRL2+ activated microglia (Fig. S10 J-L vs. M-O, S-V vs. W,Z, EE). Injection of PS19 Tau mice with LSD1 virus rescued this inflammatory response. For example, all but one (9 out of 10 analyzed) of the PS19-LSD1 inj mice had a reduction in the number of GFAP+ astrocytes (Fig. S10 G-I vs. D-F) and TRL2+ activated microglia (Fig. S10 P-R vs. M-O, AA-DD vs. W-Z, EE). Of note, the one PS19-LSD1 inj mouse where we did observe increased glial cells, similar to PS19-HA inj mice, had the lowest neuronal cell count (74% of WT-HA inj CA1 neurons). It is possible that this mouse was already undergoing neurodegeneration prior to the injection of the LSD1 overexpression virus.

**Fig. 5:**
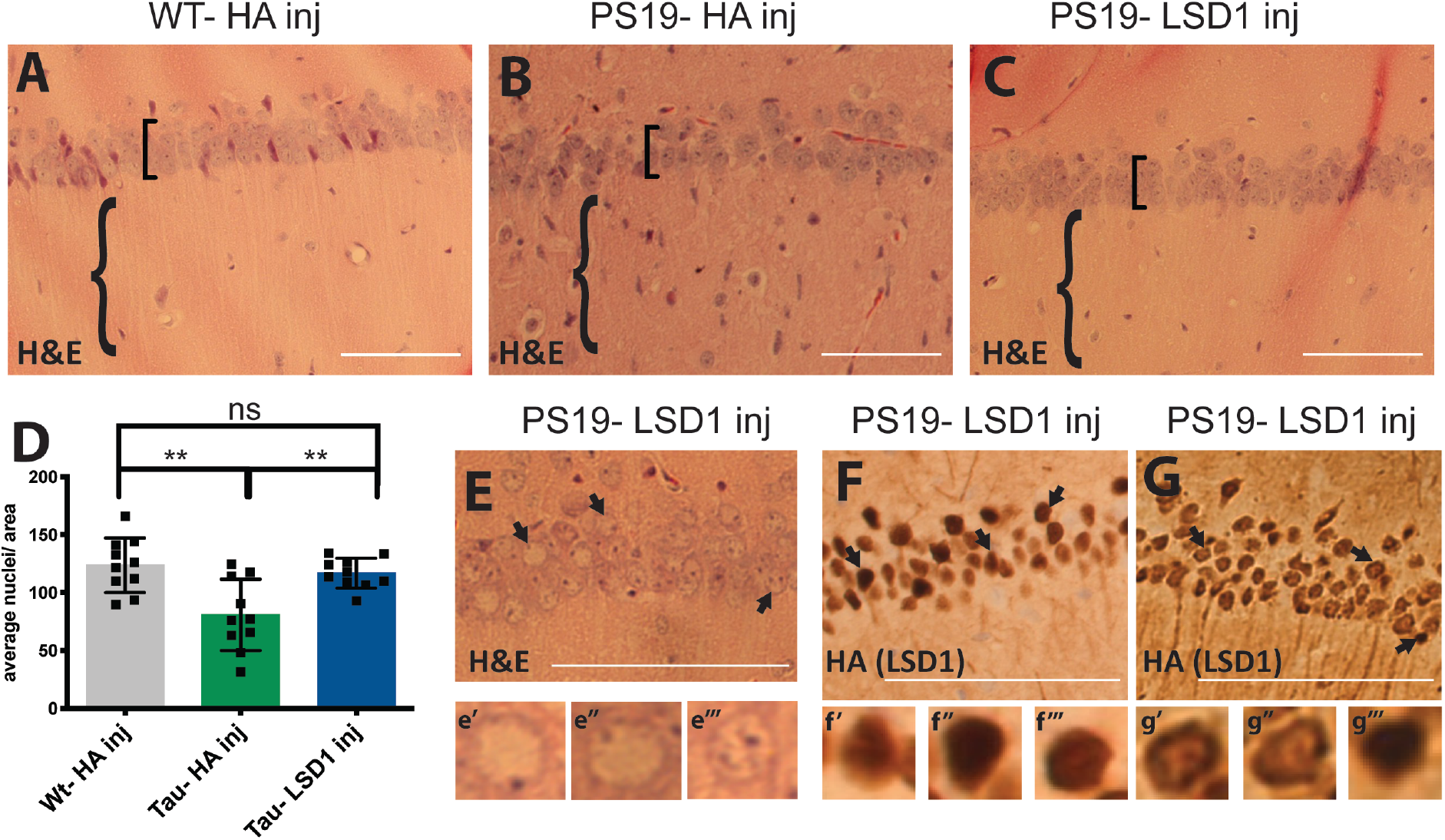
LSD1 overexpression rescues the neurodegenerative phenotype in the hippocampus of 11 month old PS19 Tau mice. **A-C,** Representative image of H&E stained CA1 region of the hippocampus of 11 month old Wild Type mice injected with HA control virus (WT-HA inj) (**A**), PS19 Tau mice injected with HA control virus (PS19-HA inj) (**B**), and PS19 Tau mice injected with *Lsd1* overexpressing virus (PS19-LSD1 inj) (**C**). Square brackets denote thickness of pyramidal layer of the CA1 of the hippocampus and curvy brackets denote hippocampal region with or without infiltrating cells. **D,** Quantification of the average number of nuclei in the pyramidal layer of the hippocampus per area per mouse from histology represented in **Fig. 5A-C**. Values are mean ± SD (*n*=10, one-way analysis of variance (ANOVA) with Tukey’s post hoc test, **p<0.01, ns=not significant). **E,** Representative H&E of PS19-LSD1 inj mouse with abnormal nuclei blebbing in the CA1 region of the hippocampus. **E’-E’’’,** High magnified image of cells denoted by arrows in **Fig. 5E** of individual nuclei that are either abnormally blebbed (**E’, E”**) or normal (**E’’’**). **F,G,** Immunohistochemistry staining of HA(LSD1) in 11 month PS19-LSD1 inj mice. HA is either localized specifically to the nucleus in all nuclei (**F**) or in only a few nuclei while it is partially sequestered in the cytoplasm in others (**G**). **F’-F’’’,** High magnified image of cells denoted by arrows in **Fig. 5F** of nuclear HA localization in individual nuclei. **G’-G’’’,** High magnified image of cells denoted by arrows in **Fig. 5G** of individual nuclei with HA(LSD1) either sequestered to the cytoplasm (**G’, G”**) or confined to the nucleus (**G’’’**). Scale bars=50μm.

Although the number of hippocampal neurons in PS19-LSD1 inj mice did not differ from WT-HA inj controls, in 6 of the 10 PS19-LSD1 inj mice we observed cells with abnormal blebbed nuclei at varying numbers throughout the hippocampus (Fig. 5E). These abnormal cells are rare in PS19-HA inj mice, which have a reduced overall number of cells in the pyramidal cell layer compared to WT-HA inj control mice. One possibility is that these abnormal cells with blebbing nuclei represent an intermediate state between a healthy neuron and a dying neuron that is prolonged by rescue via LSD1 overexpression. Interestingly, these abnormal cells also differed in the localization of HA-tagged LSD1. The four mice with normal nuclei had HA-tagged LSD1 protein localized uniformly throughout the nucleus (Fig. 5F). In contrast, the six mice with abnormal nuclear blebbing had some HA-tagged LSD1 that was mislocalized to the cytoplasm (Fig. 5G). This includes the one PS19-LSD1 inj mouse that had an elevated number of astrocytes and TRL2 positive microglia. Thus, the blebbing state correlates with when the viral produced LSD1 begins to be sequestered in the cytoplasm, similar to the endogenously produced LSD1.

To test the specificity of the rescue that we observe when LSD1 is overexpressed in the hippocampus of PS19 Tau mice, we performed RNA sequencing on the hippocampus of rescued 11-11.5 month old PS19-LSD1 inj mice versus PS19-HA inj mice. The RNA-seq analysis detected 144 significant gene expression changes PS19-HA inj mice compared to WT-HA inj control mice (Fig. S11 A,B), and 57 significant gene expression changes in PS19-LSD1 inj mice compared to WT-HA inj control mice (Fig. S11 C,D). All of the 144 significant gene expression changes induced by tau are ameliorated by overexpression of LSD1 (Fig. 6A). As a result, 116 out of the 144 significantly changed genes in the PS19 Tau-HA mice are no longer significantly changed in the rescued PS19-LSD1 inj mice. Based on this specific rescue of the differentially expressed genes induced by tau, we further compared the expression changes between PS19-LSD1 inj mice and PS19-HA inj mice genome-wide (Fig. 6B). Amongst the genome-wide expression changes in PS19-HA inj mice compared to WT-HA inj control mice, 76% were changed in the same direction (either up or down), and of these 69% were ameliorated. It should be noted, that this is the opposite of what we observe when we reduce LSD1 (Fig. 4). Thus, tau-induced gene expression changes can be specifically modulated in both directions by increasing or decreasing LSD1 levels. Importantly, the pathways identified by Gene Set Enrichment Analysis are similar between PS19-LSD1 inj mice and PS19-HA inj mice, and we fail to detect any new pathways induced by overexpression of LSD1 that could potentially account for a non-specific rescue (Fig. S11 E-H).

**Fig. 6:**
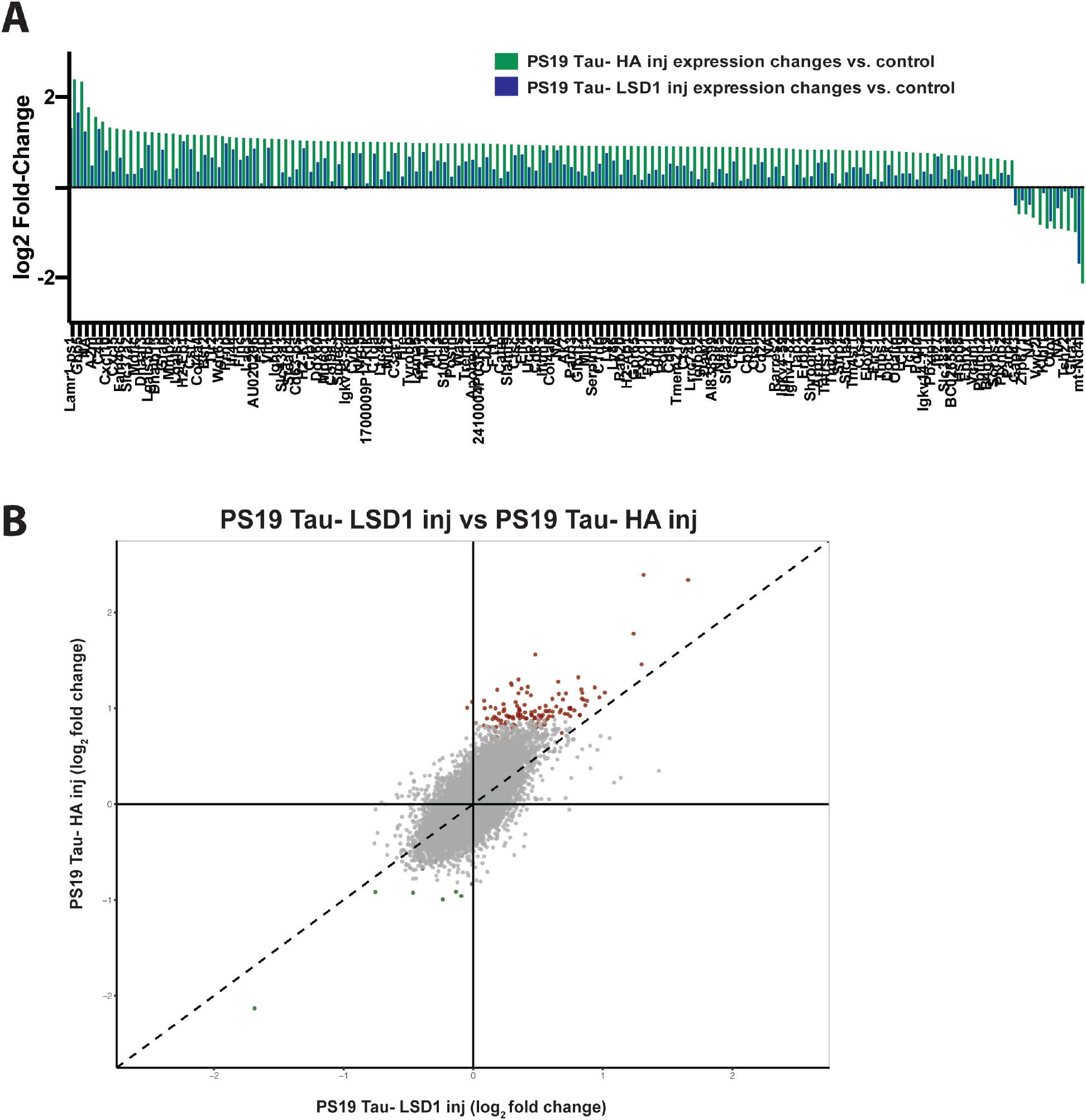
Molecular overlap between loss of LSD1 function and tauopathy. **A,** Histogram (log_2_ fold change) of the 144 genes that have significant changes in expression in the PS19 Tau-HA inj mice (green, *n*=3) and their corresponding gene expression changes in the PS19 Tau-LSD1 inj mice (blue, *n*=3). **B,** Scatter plot showing the correlation between the genome-wide log_2_ fold change in gene expression between PS19 Tau-HA inj and PS19 Tau-LSD1 inj. The most significantly changed genes in the PS19 Tau-HA inj mice are shown in red (upregulated) and green (downregulated). All other genes are shown in grey. Dotted line represents 1:1 relationship between gene expression changes in PS19 Tau-Ha inj and PS19 Tau-LSD1 inj. Rescued genes fall to the left of the dotted line in the positively correlated quadrant and to the right of the dotted line in the negatively correlated quadrant. Genes with correlated expression changes are found in the top right and bottom left quadrants, while genes that do not correlate are found in the opposite quadrants.

## DISCUSSION

In this study we investigate a potential downstream mediator of tau pathology in neurodegenerative disease. We find that modulation of the chromatin modifying enzyme LSD1 can alter neurodegeneration in a tauopathy mouse model. Previously, we showed that LSD1 colocalizes with tau pathology in the cell body of neurons in AD cases (*26*). This suggested that LSD1 might be disrupted in tauopathies such as AD, by being excluded from the nucleus. To address this directly, we utilize the PS19 tauopathy mouse model. In these mice, we find that LSD1 is sequestered in the cytoplasm, in some cases being completely depleted from the nucleus. This provides the first cytological evidence that pathological tau can prevent LSD1 from properly localizing to the nucleus in hippocampal and cortical neurons, where we have previously shown it is continuously required (*44*).

Based on the ability of pathological tau to sequester LSD1, we hypothesized that neuronal cell death may be due, at least partly, to LSD1 being sequestered in the cytoplasm and depleted from the nucleus. In this case, reducing LSD1 levels should make it easier for tau to deplete LSD1 from the nucleus, resulting in a faster progression of neurodegeneration and/or a more severe neurodegenerative phenotype. We find that reducing LSD1 in the PS19 Tau transgenic background accelerates the depletion of LSD1 from the nucleus compared to PS19 Tau mice alone. Surprisingly, in PS19;*Lsd1*^*Δ*/+^ mice we also observe a slight decrease in number of cells in the cortex with LSD1 localized completely to the cytoplasm. However, this is likely because many of the cells in mice with decreased LSD1 have died and been cleared. Consistent with this possibility, we observe less cells/area overall in PS19;*Lsd1*^*Δ*/+^ mice compared to PS19 Tau mice.

LSD1 heterozygosity alone induces only 4 significant gene expression changes and does not lead to any neurodegeneration. This suggests that any effects observed in PS19;*Lsd1*^*Δ*/+^ mice are not simply due to LSD1 haploinsufficiency. Normally, PS19 Tau mice develop paralysis and neurodegeneration, along with reduced survival. When we reduce LSD1 in the PS19 Tau mice, mice die significantly earlier, most likely due to the increased rate of paralysis. Additionally, PS19;*Lsd1*^*Δ*/+^ mice have increased neuronal cell death and clearance in the hippocampus. This suggests that pathological tau functions through LSD1 to cause neurodegeneration *in vivo* in mice.

PS19;*Lsd1*^*Δ*/+^ mice have a 31% reduction in LSD1 protein levels compared to PS19 Tau mice from birth. This reduction should theoretically make mice sensitive to LSD1 depletion from birth. However, tau pathology starts at 6-8 months in PS19 Tau mice. As a result, if *Lsd1* heterozygosity is functioning by making it easier for pathological tau to deplete LSD1 from the nucleus, we would not expect to see any exacerbation until after pathological tau is present. The exacerbation of the PS19 Tau mouse neurodegenerative phenotype does not occur until after pathological human tau was present. This suggests that the effect of *Lsd1* heterozygosity requires the presence of pathological tau, placing LSD1 downstream of tau. Consistent with LSD1 being downstream of pathological tau, we found no evidence that *Lsd1* heterozygosity affects the expression of the tau transgene, or the buildup of pathological tau in PS19 Tau mice.

To test whether the functional interaction between pathological tau and reduced LSD1 is specific, we used RNA-seq to determine whether the downstream molecular pathways altered in PS19 Tau mice are exacerbated in PS19;*Lsd1*^*Δ*/+^ mice. This analysis was performed at the time of earliest signs of neuronal distress, allowing us to assess molecular changes prior to cell death and clearance. LSD1 heterozygosity induces only 4 significant expression changes. In addition, the pathways that are affected in both PS19 Tau mice and PS19;*Lsd1*^*Δ*/+^ mice are very similar. This suggests that reducing LSD1 did not induce any additional neurodegeneration pathways. In contrast, when LSD1 is reduced in PS19 Tau mice, the genome-wide expression changes induced by pathological tau are specifically exacerbated. This suggests that the functional interaction that we observe between pathological tau and reduced LSD1 is occurring through the tau pathway.

Our previous data implicated LSD1 in the tau-mediated neurodegeneration pathway. Utilizing the PS19 Tau mouse model, we now show a functional interaction between pathological tau and LSD1. Importantly, because the tau mutation utilized in PS19 Tau mice comes from an FTD patient, the functional interaction that we detected may also be relevant to FTD and other taupathies, such progressive supranuclear palsy, corticobasal degeneration and chronic traumatic encephalopathy. However, because PS19 Tau mice do not have Aβ plaque accumulation, this functional interaction would not be relevant to dysfunction induced directly by pathological Aβ.

Based on the data we uncovered showing a specific functional interaction between pathological tau and LSD1, we propose the following model (Movie S4): in healthy hippocampal and cortical neurons, LSD1 is translated in the cytoplasm and transported into the nucleus where it is continuously required to repress inappropriate transcription. In tauopathy, pathological tau accumulates in the cytoplasm blocking LSD1 from nuclear import. This interferes with the continuous requirement for LSD1, resulting in neuronal cell death. It is possible that pathological tau inhibits LSD1 function indirectly, by interfering with some other process that subsequently disrupts LSD1 function. However, based on our observations that LSD1 mislocalizes to cytoplasmic pathological tau in PS19 Tau mice and human AD cases (*44*), we favor a model where pathological tau directly masks LSD1 from interacting with the nuclear import complex. Regardless of whether the interaction is direct or indirect, it should be noted that breakdown of the nuclear pore, which has been observed in human AD cases (*45–47*), would potentially exacerbate the model that we propose.

This model makes a direct prediction: if tau is predominantly functioning through LSD1, then increasing the levels of LSD1 should rescue the tau-induced neurodegenerative phenotype. To address this, we overexpressed LSD1 in the hippocampal neurons of PS19 Tau mice. Overexpression of LSD1 specifically in hippocampal neurons rescues the neuronal cell death and limits the inflammatory response. This rescue is neuronal specific, suggesting that the functional interaction between LSD1 and tau is occurring in neurons. In addition, this rescue occurs despite there being no effect on tau aggregation. This negates the possibility that the tau transgene is simply not functioning when LSD1 is overexpressed. The ability of LSD1 overexpression to overcome tau-mediated neurodegeneration in the presence of pathological tau aggregates, provides further evidence that pathological tau is functioning through the inhibition of LSD1.

*A priori*, we would not expect the gene expression changes induced by overexpression of LSD1 to be related to the gene expression changes induced by pathological tau. However, the data presented here, along with our previous work (*44*), raise the possibility that pathological tau may be causing neurodegeneration by sequestering and inhibiting tau. If this is the case, we would expect that overexpression of LSD1, even after pathological tau is present, would specifically counteract tau-induced expression changes. Strikingly, we find that overexpression of LSD1 ameliorates tau-induced gene expression changes genome-wide. This suggests that the inhibition of LSD1 may be the critical mediator of neurodegeneration caused by pathological tau.

Overexpressing LSD1 should not prevent it from being sequestered. Rather overexpressing LSD1 should make it more difficult for pathological tau to sequester all of the LSD1 protein, allowing some LSD1 to be transported to the nucleus. Thus, overexpressing LSD1 would be expected to temporarily rescue the ability of pathological tau to kill neurons. Consistent with this, LSD1 overexpression delays the effect of pathological tau rather than permanently rescuing. In 60% of the mice, the surviving neurons have abnormal morphology, and the overexpressed version of LSD1 is also sequestered. The observation that neurons fail to maintain their morphology when the overexpressed LSD1 begins to be sequestered in the cytoplasm provides further support for the model that tau mediates neurodegeneration through the sequestration of LSD1. Nevertheless, our data suggest that overexpression of LSD1 cannot permanently overcome pathological tau. To permanently overcome pathological tau, it would likely be necessary to permanently disrupt the interaction between pathological tau and LSD1. This work is currently ongoing in the lab. Overall, our data establish LSD1 as a major downstream effector of tau-induced neurodegeneration. Based on these data, we propose that the LSD1 pathway is a potential late stage target for intervention in tauopathies, such as AD.

## MATERIALS AND METHODS

The mouse strains are described in SI Materials and Methods. All relevant techniques are also included there. These include mouse tissue fixation, histology and histological studies, immunohistochemistry and immunofluorescence, protein quantification, quantitative analysis of paralysis, MRI of brain atrophy, RNA sequencing, RNA sequencing analysis, stereotaxic surgery and viral infusion, and statistical analysis.

## ACKNOWLEDGMENTS

We thank M. Rosenfeld (U.C.S.D) for providing *Lsd1* deletion mice and D. Castrillon for providing the *Vasa-Cre* mice; N. Seyfried, R. Betarbet, Mr. Gearing, from the Emory ADRC (P50 AG025688), NINDS Emory Neuroscience Core Facilities (P30NS055077) for tissue processing and providing PHF1 antibody, J. Alcudia of the Stanford Gene Vector and Virus Core for the help with virus generation and production, the Georgia Genomics and Bioinformatics Core, which provided the RNA library preparation and sequencing service, J. Schroeder from Emory Rodent Behavioral Core for help with Rotatord and Grid Performance assays supported by the Emory Neuroscience NINDS Core Facilities (P30NS055077) with further support provided by the Georgia Clinical & Translational Science Alliance of the National Institutes of Health under Award Number UL1TR002378; and J. Park from the Emory Center for Systems Imagining for preforming MRI supported by the National Center for Advancing Translational Sciences of the National Institutes of Health under Award Number UL1TR000454. We would also like to thank M.J. Rowley and V. Corces for assistance on RNA sequencing analysis, K. Porter-Stransky and D. Weinshenker for providing PS19 Tau mouse and teaching stereotaxic surgery procedure, L.R. Lym and Dr. Lerit for assistance with confocal imagining; D. Weinshenker, A. Levey, C. Bean and T. Caspary for comments on the manuscript and assistance throughout. Additionally the authors would like to thank fellow Katz lab members for assistance experimentally and intellectually.

## Data and materials availability

FastQ files for the RNA sequencing experiments are bring deposited in the GEO and will be available upon publication. Results from DESEQ2 are available in Data S1. Correspondence and requests for materials should be addressed to D.J.K.

