## Supplementary material for "The inhibition of LSD1 via sequestration contributes to tau-mediated neurodegeneration": materials and methods

All mouse work, including surgical procedures, were approved by and conducted in accordance with the Emory University Institutional Animal Care and Use Committee.

Animals

To generate tauopathy mice that are heterozygous for *Lsd1*, PS19 Tau P301S mice (Jackson Laboratory stock nos. 008169, generated by the Lee lab (*28*)) were crossed with mice that are heterozygous for *Lsd1* (*Lsd1*$^{\Delta/+}$). *Lsd1*$^{\Delta/+}$ mice were generated by crossing *Lsd1*$^{fl/+}$ mice, generated in the Rosenfeld lab (*27*), with *Vasa-Cre* transgenic mice(*31*). Once the deletion allele passes through the germline, *Lsd1* is heterozygous throughout the animal.

Mouse tissue fixation

Mice were given a lethal dose of isoflurane via inhalation, then transcardially perfused with ice cold 4.0% paraformaldehyde in 0.1M phosphate buffer. Brain and spinal cord were dissected and post fixed in cold paraformaldehyde solution for 2 hours. Brain weights and sizes were taken from mice that were euthanized by cervical dislocation. Brain was dissected, immediately weighed, imaged, and fixed in cold 4.0% paraformaldehyde in 0.1M phosphate buffer overnight. In all cases, tissues were transferred to cold PBS, then serially dehydrated and embedded in paraffin and serially sectioned into 8µm coronal sections.

Histology and histological studies

Hematoxylin and eosin staining was performed according to standard procedures. Briefly, sections were dewaxed with xylenes and serial ethanol dilutions then stained with Eosin using the Richard-Allan Scientific Signature Series Eosin-Y package (ThermoScientific). To derive unbiased estimates of neuronal loss in the hippocampus, the number of primordial neurons in CA1 and CA3 (corresponding approximately to bregma coordinates -2.0 mm and -3.0 mm) were counted from 2 randomly selected regions in the field of a Zeiss Axiophot ocular graticule grid and measured manually using digital micrographs of H&E-stained preparations. Investigators were blinded to the genotype or treatment.

Immunohistochemistry and immunofluorescence

Sections were dewaxed with xylenes and serial ethanol dilutions, then treated with 3% hydrogen peroxide at 40°C for 5 minutes to quench endogenous peroxidase activity, blocked in 2% serum at 40°C for 15 minutes, and incubated with primary Ab (Table S1) overnight at 4°C. Slides were washed, then incubated with biotinylated secondary Ab (Biotinylated Goat α Rabbit, 1:200, Vector Labs BA-1000 or Biotinylated Goat α Mouse, 1:200, Vector Labs BA-9200) at 37°C for 30 minutes. Signal amplification was then carried out by incubating at 37°C for 1 hour with Vector Labs Elite ABC reagent (PK-6200). Slides were developed with DAB for 1-5 minutes, counterstained with hematoxylin for 1 minutes, and coverslipped. For immunofluorescence, dewaxed sections were first rinsed with TBS. Antigen retrieval was performed by microwaving at 10% power 2X for 5 minutes in 0.01M sodium citrate. Slides were then cooled, washed with TBS, permeabilized in 0.5% Triton X-100 for 20 minutes, followed by blocking in 10% goat serum 20 minutes. Primary Abs (Table S1) were incubated overnight at 4°C. Slides were then washed and incubated in secondary Abs (Invitrogen A1 1001 and Invitrogen A11012) for 1 hour at room temperature, followed by TBS washes, counterstained with DAPI, and then coverslipped. For the assessment of tau accumulation, six random sections (sampling from CA1, CA3, and cerebral cortex) per sample were manually counted using digital micrographs of AT8 stained preparations in the field of a Zeiss Axiophot ocular graticule grid. Investigators were blinded to the genotype or treatment. Imagining for immunofluorescence of LSD1 staining was performed on a spinning-disk confocal Nikon-Tie controlled with the software NIS Elements

(Nikon). Imaging for all other immunofluorescence staining was performed on an Eclipse Ti2 inverted microscope (Nikon, Toyko, Japan) controlled with the software NIS Elements (Nikon). Image J software ((NIH, http://imagej.nih.gov/ij/) was used for viewing all images.

Protein Quantification

Protein levels were determined by homogenizing brains in 1 ml/g of tissue in ice-cold lysis buffer (150mM NaCl, 1% Triton X-100, 0.5% Na-deoxycholate, 1% SDS, 50mM Tris, pH8.0) in a dounce homogenizer, followed by end-over-end spin at 4°C for 2 hours, and centrifugation at 20,000 x g for 20 minutes at 4°C. Protein concentrations were determined following standard BCA protocol (Pierce BCA Protein Assay Kit). Equal amounts of protein for each sample were loaded and run on a 12% SDS-PAGE gel, transferred (Semi-dry transfer using BIO RAD Trans-Blot Turbo Transfer System), blocked in 5% BSA, and probed with primary Ab (Table S1) over night at 4°C. Blots were rinsed and stained with HRP-conjugated secondary Ab, and detected by chemiluminescence using ChemiDoc MP Imaging System (BIO RAD). Protein levels were normalized using total protein calculated using BIO RAD ImageLab software.

Quantitative analysis of paralysis

We performed experiments on PS19 Tau, *Lsd1*$^{\Delta/+}$, PS19;*Lsd1*$^{\Delta/+}$ mice at 6, 8, and 10 months. For the rotarod experiments, mice were given two practice trials and then placed on the rotating cylinder at 4rpm. Rotational speed then gradually increased over a 5-minute test session up to a maximum rotational speed of 40rpm. Latency to fall off of the accelerating rotarod was used as the dependent variable. We calculated the latency to fall, maximum speed in rotations per minute, and distance traveled. For grid performance, mice were placed on a horizontal grid that was then inverted so mice are hanging upside down by their paws. Mice were videotaped for 10 seconds, and then scored for forepaw and back paw distance traveled. Mice that could not hold onto grid for 10 seconds were censored. Investigators were blinded to the genotypes for both experiments.

MRI of brain atrophy

MRI studies were conducted on PS19 Tau, *Lsd1*$^{\Delta/+}$, PS19;*Lsd1*$^{\Delta/+}$ mice at 6 months and 10 months (*n*=3/genotype). Mice were anesthetized with isoflurane, and monitored for heart rate and temperature changes while anesthetized. MRI measurements were performed using a 9.4 T/20 cm horizontal bore Bruker magnet, interfaced to an AVANCE console (Bruker, Billerica, MA, USA). A two-coil actively decoupled imaging set-up was used (a 2 cm diameter surface coil for reception and a 7.2 cm diameter volume coil for transmission). Axial T2-weighted images were acquired with a RARE (Rapid Acquisition with Refocused Echos) sequence. Its imaging parameters were as follows: TR = 3000 ms, Eff.TE = 64 ms, RARE factor = 4, field of view (FOV) = 23.04 × 23.04 mm2, matrix = 192 × 192, Avg = 4, slice thickness (thk) = 0.6 mm, number of slice(NSL)=20. Specific emphasis was placed on the neocortex and hippocampus in the coronal images (1.0 – 4.0 mm posterior to the bregma).

RNA sequencing

9 month old *Lsd1*$^{+/+}$, *Lsd1*$^{\Delta/+}$, PS19 Tau, and PS19;*Lsd1*$^{\Delta/+}$ littermates (n=2 mice/genotype) were euthanized by cervical dislocation, hippocampi were dissected and snap frozen with liquid nitrogen in 1mL Trizol, and stored at -80°C. For RNA isolation, samples were thawed at 37°C then kept on ice prior to homogenization with Polytron homogenizer with a 5 second pulse. After a 5 minute incubation at room temperature, one tenth the sample volume of 1-bromo-3chloropropane was added, mixed by inversion and incubated for 3 minutes at room temperature. Samples were then centrifuged at 13,000 X g for 15 minutes at 4°C to separate the aqueous and

organic layers. As much of the aqueous layer was recovered as possible, then RNA was precipitated with isoproponal. Pellets were then washed with 75% ethanol and resuspended in 50μL of dionized water. RNA library preparation and sequencing were performed by HudsonAlpha Genomic Services Lab. RNA was Poly(A) selected and 300bp size selected. Libraries were sequenced for 25 million 50bp paired end reads. For analysis of PS19 Tau mice injected with virus, the hippocampus was processed as above. RNA library preparation and sequencing were performed by Georgia Genomics and Bioinformatics Core. RNA was Poly(A) selected and 300bp selected. Libraries were sequenced generating ~36 million reads per sample, 75bp paired end reads.

RNA sequencing analysis

The sequencing data were uploaded to the Galaxy web platform, and we used the public server at *usegalaxy.org* to analyze the data (48, 49). FASTQ files were quality assessed using FASTQC (v.0.11.7), trimmed using Trimmomatic (v.0.36.5) and minimum QC score of 20 and minimum read length of 36bp. Paired-end reads were subsequently mapped to the GRCm38 genome using HISAT2 (v.2.1.0). Unmapped, unpaired and multiply mapped reads were removed using Filter SAM or BAM (v.1.1.2). Assignment of transcripts to GRCm38 genomic features was performed using Featurecounts (v.1.6.0.6) and the Ensembl GRCm38.93 gtf file. Differentially expressed transcripts were determined using DESEQ2 (v.2.11.40.2) *(49)*. For all datasets, a cutoff of adjusted p-value < 0.3 and abs ($\log_2$ fold change) > 0.58 was applied. TPM values were calculated from raw data obtained from Featurecounts output. Subsequent downstream analysis was performed using R and normalized counts and adjusted P-values from DESEQ2 (v.2.11.40.2). Heatmaps were produced and hierarchical clustering was done using the gplots package (v. 3.0.1) and normalized counts *(50)*. Volcano plots were produced using the enhanced volcano package (v.0.99.16) and adjusted p-values *(51)*. Additionally, Gene Set Enrichment Analysis (Pre-ranked list) was performed using the online platform WebGestalt *(52-55)*. Custom R-scripts available upon request.

Stereotaxic surgery and viral infusion

All surgical procedures were approved by and conducted in accordance with the Emory University Institutional Animal Care and Use Committee. Mice were anesthetized with isoflurane (3% induction, 1-2% maintenance) and administered the analgesic meloxicam (5 mg/kg). Using a Stoeling Quintessential Stereotaxic Injector pump and Hamilton syringe, mice were injected with either the AAV-DJ-LSD1- HA virus or the control AAV-DJ- HA virus into both hippocampi. Each virus was injected into the rostral (AP: -2.5, ML:± 2.2, DV: -1.6, relative to bregma) and caudal (AP: -3.1, ML:± 3.0, DV: -3.5) hippocampus of both hemispheres (four injection sites total). Infusion volumes were 0.5 µL per injection site, administered at a rate of 0.15 µL/min. Following surgery, mice were monitored daily for the duration of the experiment. Brains were extracted 3 months post-surgery which allows sufficient time for viral expression. Injection accuracy was confirmed by HA positive staining, and those mice where staining was outside the hippocampus or that did not fully reach hippocampus were censored.

Statistical analysis

Data are expressed as the either the mean± SD (SEM for LSD1 localization and paralysis quantification), or box-and-whisker plots where the box plot edges are $25^{th}$ and $75^{th}$ percentile, central line is the median, and whiskers are maximum and minimum. Because the data were normally distributed, parametric tests were used to analyze the data in GraphPad Prism version 7 (GraphPad software, San Diego, CA). Data were analyzed via one-way or two-way ANOVA with post hoc Tukey's test for multiple comparisons when appropriate. The survival curve was analyzed using the Log-rank Mantle-Cox test. Significance was set at $P<0.05$ and two-tailed variants of tests were used throughout. Each specific test and number of independent experimental trials (*n*) are described in the figure legends coordinating with data shown.
