## supplemental figure 1 for "The inhibition of LSD1 via sequestration contributes to tau-mediated neurodegeneration"

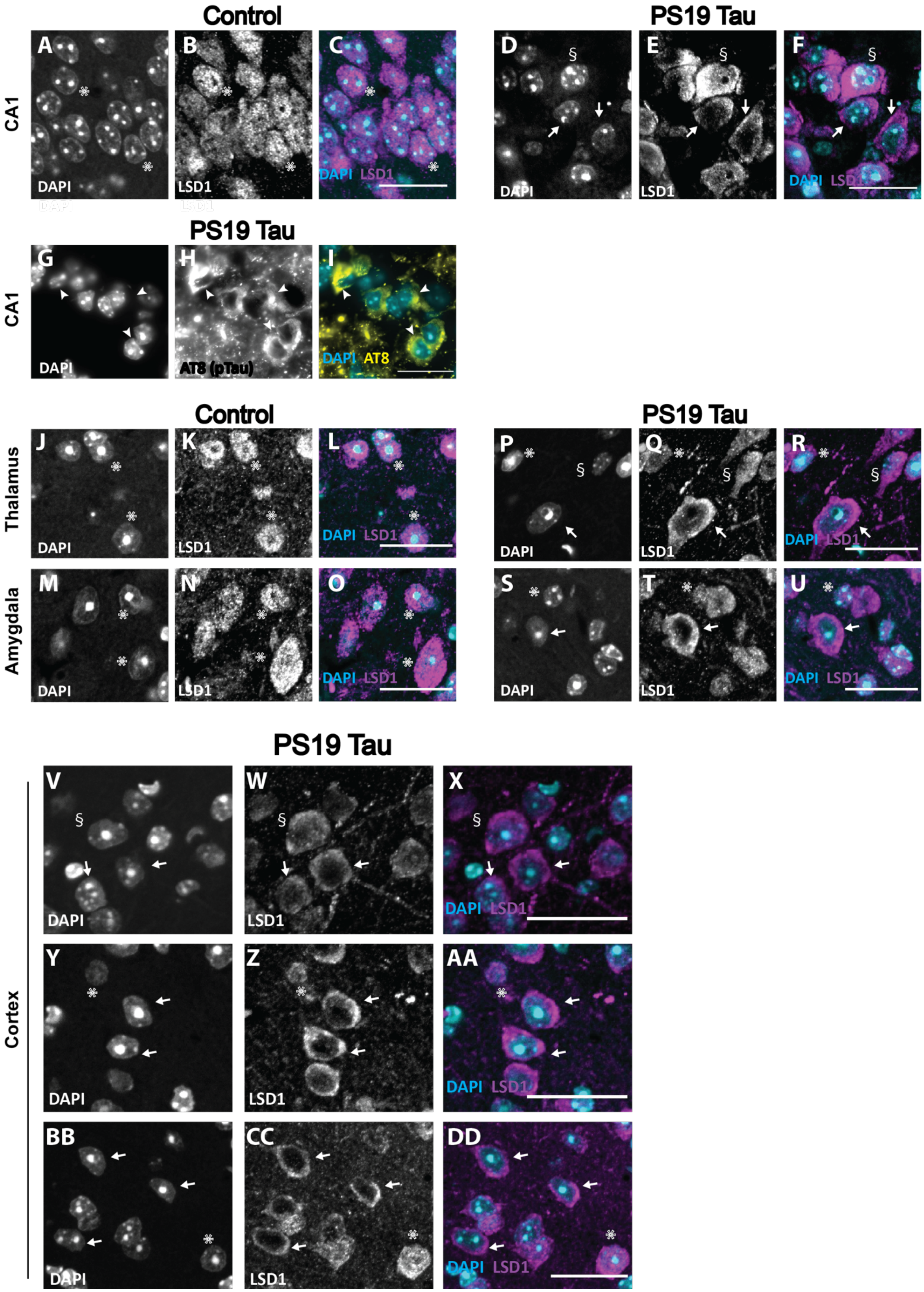

Control
PS19 Tau
CA1
A
B
C
D
E
F
DAPI
LSD1
DAPI LSD1
§
PS19 Tau
G
H
I
DAPI
AT8 (pTau)
DAPI AT8
Control
PS19 Tau
Thalamus
Amygdala
J
K
L
M
N
O
P
Q
R
S
T
U
PS19 Tau
Cortex
V
W
X
Y
Z
AA
BB
CC
DD

**Fig. S1: Sequestration of LSD1 in PS19 Tau mice.** **A**-**F**, Representative immunofluorescence showing DAPI (**A,D**), LSD1 (**B,E**), and merged (**C,F**) images in the CA1 region of the hippocampus in 12 month old Wild Type (**A-C**) and PS19 mice (**D-F**). **G-I**, Representative immunofluorescence showing DAPI (**G**), AT8 positive hyper-phosphorylated tau (**H**), and merged (**I**) in the CA1 region of the hippocampus of 12 month old PS19 Tau mice showing hyper-phosphorylated tau accumulation in the cytoplasm of the cell bodies. Arrowheads denote hyper-phosphorylated tau. **J-U,** Representative immunofluorescence showing DAPI (**J,M,P,S**), LSD1 (**K,N,Q,T**), and merged (**L,O,R,U**) images in the thalamus (**J-L,P-R**) and amygdala (**M-O,S-U**). In 12 month old control Wild Type mice (**J-O**), LSD1 is localized specifically to the DAPI positive nuclei, but in 12 month old PS19 Tau mice (**P-U**) LSD1 is localized outside of the nucleus. **V-DD,** Additional examples of immunofluorescence showing DAPI (**V,Y,BB**), LSD1 (**W,Z,CC**), and merged (**X,AA,DD**) of the cerebral cortex of 12 month old PS19 Tau mice. Arrows denote cells where LSD1 is localized outside of the nucleus, asterisks denote LSD1 localized specifically to the nucleus, and § denotes cells where LSD1 is both nuclear and cytoplasmic. *n*=7 mice analyzed (images representative of 6 of the 7 mice analyzed). Scale bars=25μm.
