## supplemental figures 2-5 for "The inhibition of LSD1 via sequestration contributes to tau-mediated neurodegeneration"

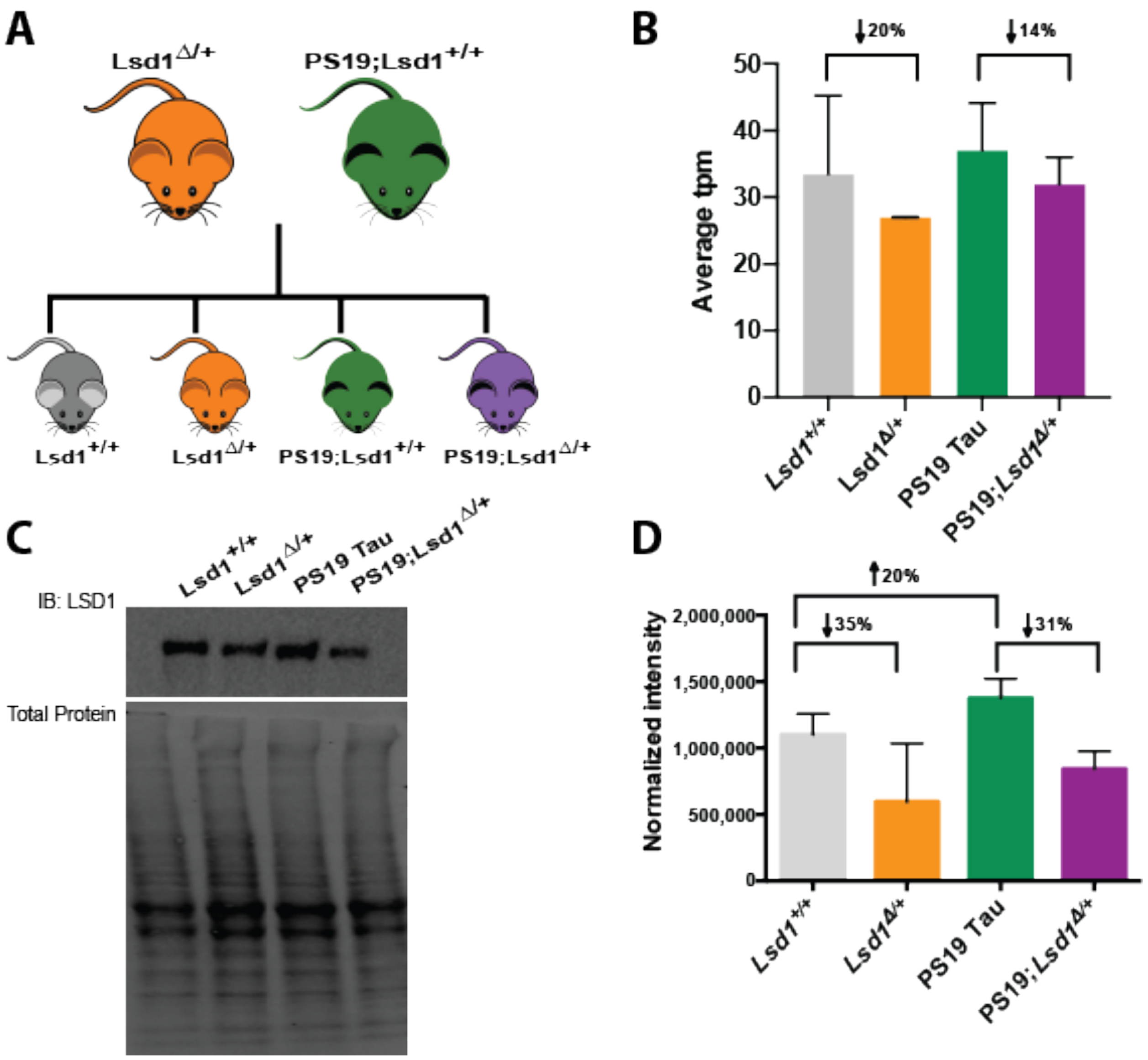


**Fig. S2: Generation of PS19 Tau mice with reduced levels of LSD1. A,** PS19 Tau mice carrying the P301S human tau transgene that are wild-type for *Lsd1* were crossed with *Lsd1* heterozygous mice. These crosses generated four genotypes: Wild Type mice (*Lsd1*$^{+/+}$, grey), *Lsd1* heterozygous mice (*Lsd1*$^{\Delta/+}$, orange), PS19 Tau mice that are wild-type for *Lsd1* (PS19;*Lsd1*$^{+/+}$ referred to as PS19 Tau, green), and PS19 Tau mice that are heterozygous for *Lsd1* (PS19;*Lsd1*$^{\Delta/+}$, purple). Colors designated here are maintained throughout all figures. **B,** Average transcripts per million (tpm) from RNA-sequencing of *Lsd1* expression in the hippocampus of *Lsd1*$^{+/+}$(*n*=4), *Lsd1*$^{\Delta/+}$(*n*=2), PS19 Tau (*n*=4), and PS19;*Lsd1*$^{\Delta/+}$ (*n*=4) mice. *Lsd1*$^{\Delta/+}$ mice had a 20% reduction in expression compared to *Lsd1*$^{+/+}$ mice, and PS19;*Lsd1*$^{\Delta/+}$ had a 14% reduction in expression compared to PS19 Tau mice. Values are mean ± SD, one-way analysis of variance (ANOVA) **P<0.01, ***P<0.005). **C,** Representative image of protein levels in the brain of *Lsd1*$^{+/+}$, *Lsd1*$^{\Delta/+}$, PS19 Tau, and PS19;*Lsd1*$^{\Delta/+}$ mice from LSD1 immunoblot and corresponding total protein blot. **D,** Quantification of immunoblot for LSD1 normalized to total protein loaded per sample as represented in **C**. Compared to *Lsd1*$^{+/+}$ mice, *Lsd1*$^{\Delta/+}$ mice had a 35% reduction and PS19 Tau mice had 20% increase in LSD1 protein levels. PS19; *Lsd1*$^{\Delta/+}$ mice had a 31% reduction in LSD1 protein level compared to PS19 Tau mice. Values are mean ± SD (*n*=3, one-way analysis of variance (ANOVA)).

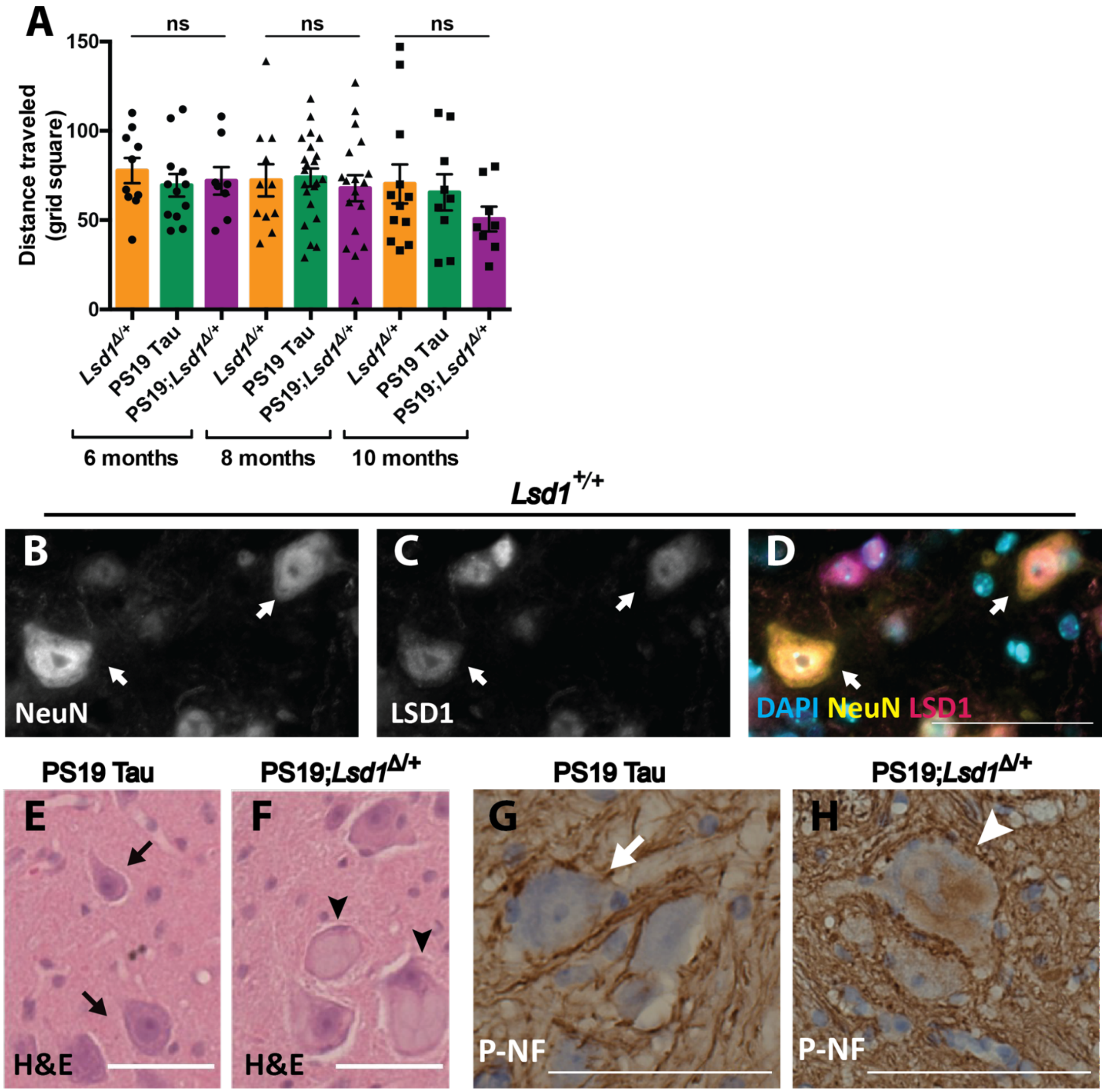


**Fig. S3: Reduction of *Lsd1* affects spinal cord in PS19 Tau mice. A,** Grid performance test measuring the distance traveled (grid squares traversed) with both forelimbs and hindlimbs in 6, 8, and 10 month old mice. *Lsd1*$^{\Delta/+}$ (orange, *n*=10,11,12), PS19 Tau (green, *n*=12,22,9), and PS19;*Lsd1*$^{\Delta/+}$ (purple, *n*=8,18,8). Values are mean ± SEM (two-way analysis of variance (ANOVA) with Tukey's post hoc test. ns=not significant). **B-D,** Immunofluorescence staining of NeuN (**B**), LSD1 (**C**), and merged with DAPI (**D**) in spinal cord motor neurons of 12 month old *Lsd1*$^{+/+}$ control mice. **E,F,** Representative image of hematoxylin and eosin (H&E) staining of motor neurons in 12 month old PS19 Tau mice (**E**) and PS19;*Lsd1*$^{\Delta/+}$ (**F**) littermates. **G,H** Representative image of immunohistochemistry staining for phospho-nuerofilament (brown) counterstained with DAPI (blue) in the motor neurons of 12 month old PS19 Tau mice (**G**) and PS19;*Lsd1*$^{\Delta/+}$ (**H**) littermates. Arrows denote healthy motor neurons. Arrowheads denote abnormal motor neurons. Scale bars=50µm.

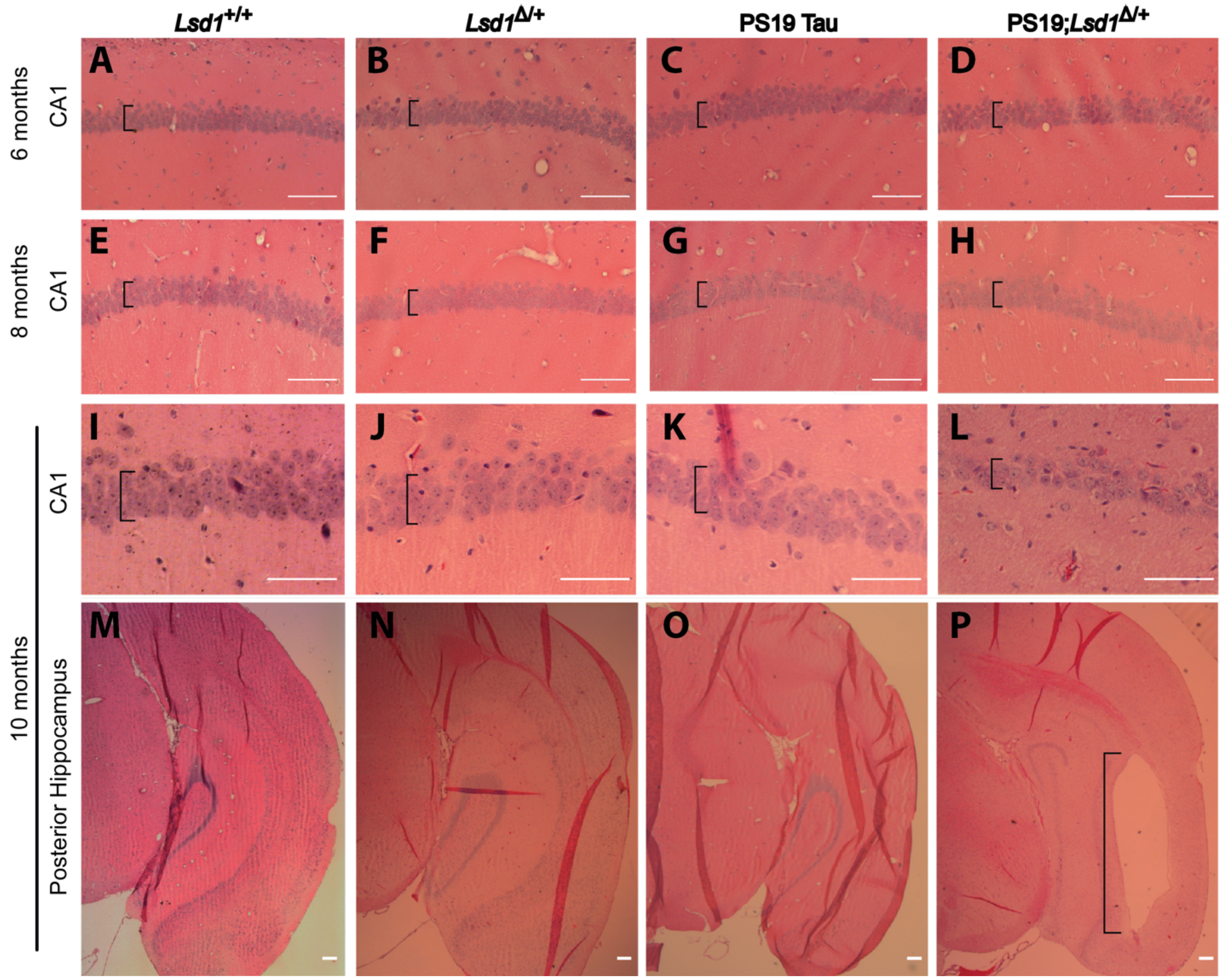


**Fig. S4: There is no exacerbation of neurodegeneration in PS19 Tau mice with reduced *Lsd*1 until 10 months of age. A-P,** Representative image of H&E staining of *Lsd1*$^{+/+}$ (**A**,**E**,**I**,**M**), *Lsd1*$^{\Delta/+}$ (**B**,**F**,**J**,**N**), PS19 Tau (**C**,**G**,**K**,**O**), and PS19;*Lsd1*$^{\Delta/+}$ (**D**,**H**,**L**,**P**) littermates at 6 months (**A-D**), 8 months (**E-H**) and 10 months (**I-P**) in the CA1 (**A-L**) and posterior hippocampus (**M-P**). Brackets denote thickness of pyramidal layer of the CA1 (**A-L**), and region of cell clearance in posterior hippocampus (**P**). Scale bars=50µm.

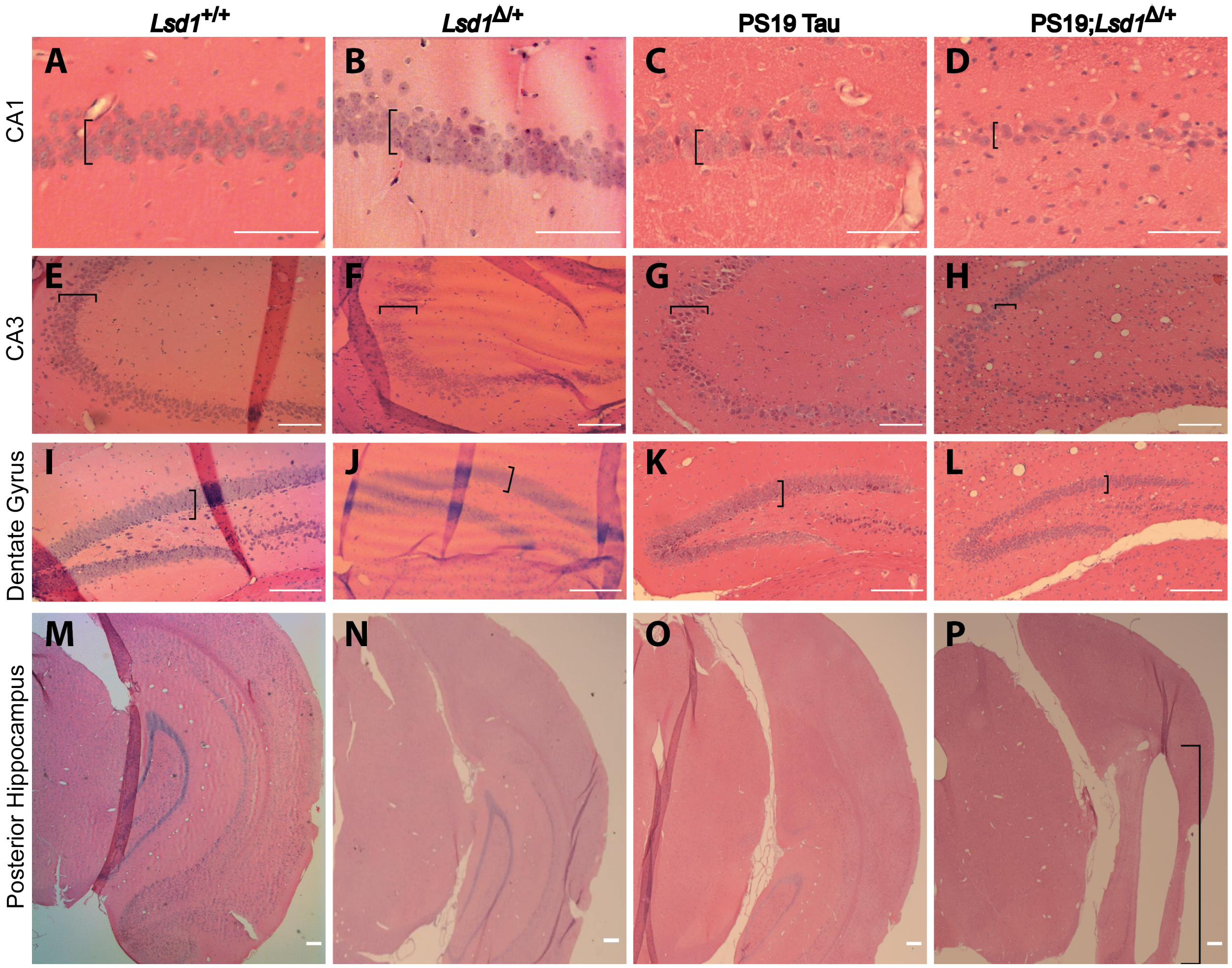


**Fig. S5: Increased neurodegeneration throughout the hippocampus and cortex of 12 month old mice. A-P,** H&E staining of 12 month old *Lsd1*$^{+/+}$ (**A,E,I,M**), *Lsd1*$^{\Delta/+}$ (**B,F,J,N**), PS19 Tau (**C,G,K,O**), and PS19;*Lsd1*$^{\Delta/+}$ (**D,H,L,P**) littermates in the CA1 (**A-D**) and CA3 (**E-H**) regions of the hippocampus, the dentate gyrus (**I-L**), and the posterior hippocampus (**M-P**). Brackets denote thickness of pyramidal layer of the CA1 (**A-D**), CA3 (**E-H**), the granule cell layer of the Dente Gyrus (**I-L**), and region of cell clearance in posterior hippocampus (**P**). Scale bars=50μm.
