## supplemental figures 6-7 for "The inhibition of LSD1 via sequestration contributes to tau-mediated neurodegeneration"

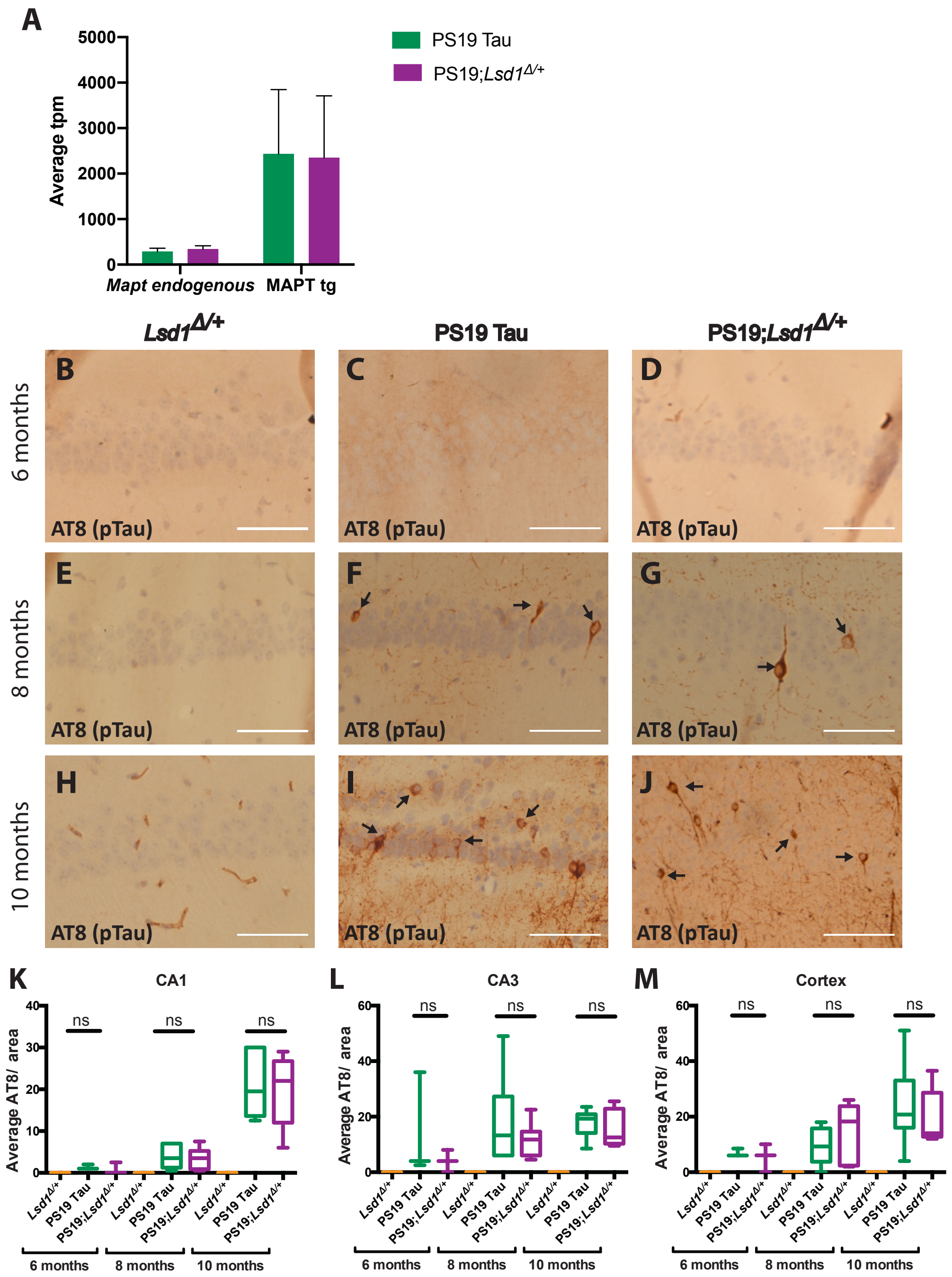


**Fig. S6: Reduction of *Lsd1* does not affect AT8 positive tau pathology. A,** Average transcripts per million (tpm) from RNA sequencing of endogenous MAPT and the expression of the human P301S MAPT transgene in the hippocampus of PS19 Tau, and PS19; *Lsd1*$^{\Delta/+}$ mice. Values are mean ± SD (*n*=4). **B-J,** Representative image of immunohistochemistry staining of phosphorylated tau (AT8 antibody) of the CA1 region of the hippocampus in *Lsd1*$^{\Delta/+}$ (**B,E,H**) , PS19 Tau (**C,F,I**), and PS19;*Lsd1*$^{\Delta/+}$ (**D,G,J**) littermates at 6 months (**B-D**), 8 months (**E-G**), and 10 months (**H-J**). Arrows denote AT8 positive immunoreactivity. Scale bars=50µm. **K-M,** Quantification of the average AT8 positive tau immunoreactivity per area from histology represented in **b-j** in the CA1 (**k**) and CA3 (**l**) regions of the hippocampus, and the cerebral cortex (**m**) (6 months *n*=3, 8 months *n*=6, and 10 months *n*=6, box plot edges are 25$^{th}$ and 75$^{th}$ percentile, central line is the median, and whiskers are max and min). For all graphs: one-way analysis of variance (ANOVA) with Tukey's post hoc test (two-sided), ns=not significant.

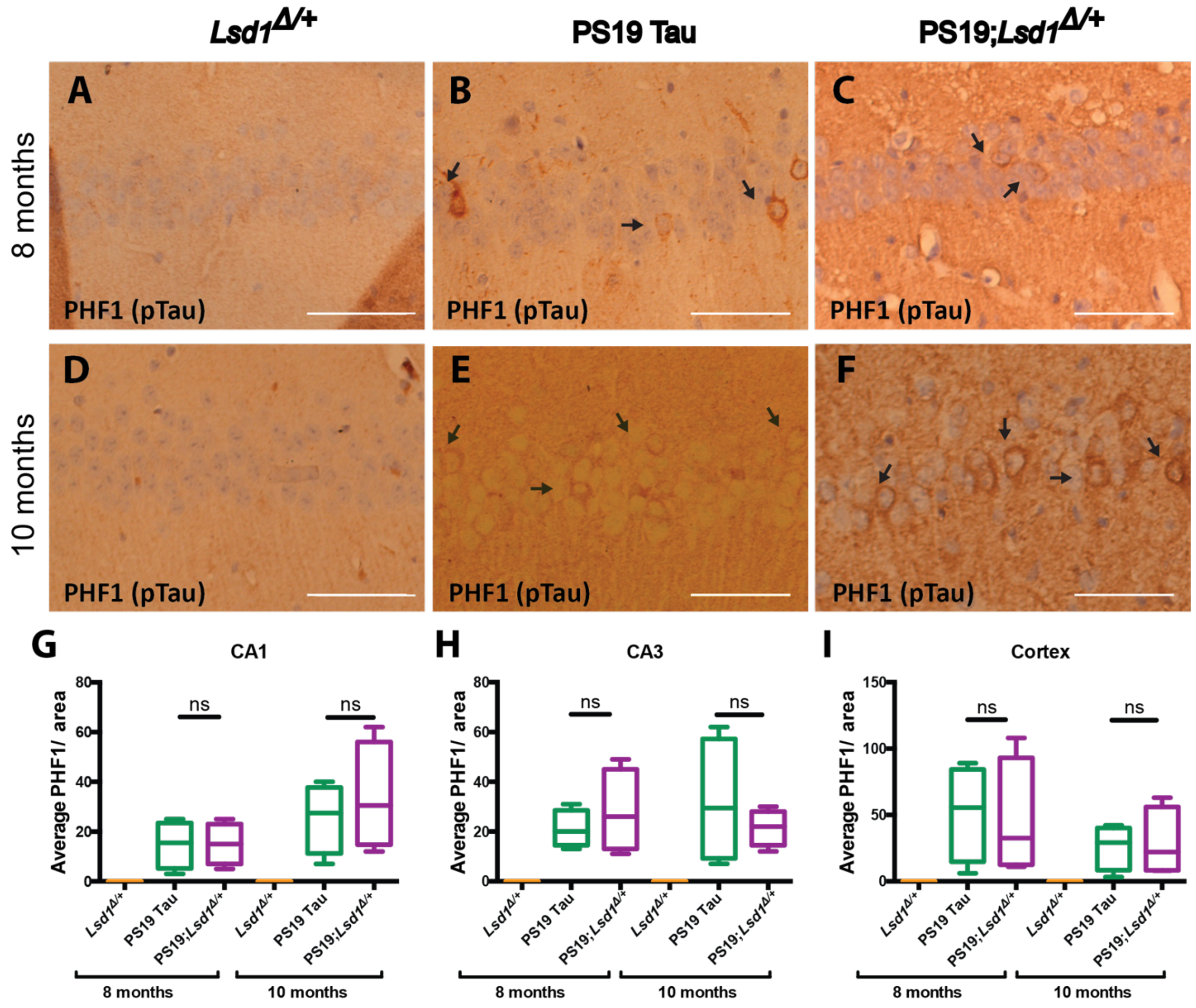


**Fig. S7: Reduction of *Lsd1* does not affect PHF1 positive tau pathology. A-F,** Representative image of immunohistochemistry staining of PHF1 in the CA1 region of the hippocampus in *Lsd1*$^{\Delta/+}$ (**A,D**), PS19 Tau (**B.E**), and PS19;*Lsd1*$^{\Delta/+}$ (**C,F**) littermates at 8 months (**A-C**) and 10 months (**D-F**). Arrows denote PHF1 positive immunoreactivity. Scale bars=50μm. **G-I,** Quantification of average PHF1 positive tau immunoreactivity per area from histology represented in **a-f** in the CA1 (**G**) and CA3 (**H**) regions of the hippocampus, and the cerebral cortex (**I**) (*n*=4 box plot edges are 25$^{th}$ and 75$^{th}$ percentile, central line is the median, and whiskers are max and min). For all graphs: one-way analysis of variance (ANOVA) with Tukey's post hoc test (two-sided), ns=not significant.
