## supplemental figure 8 for "The inhibition of LSD1 via sequestration contributes to tau-mediated neurodegeneration"

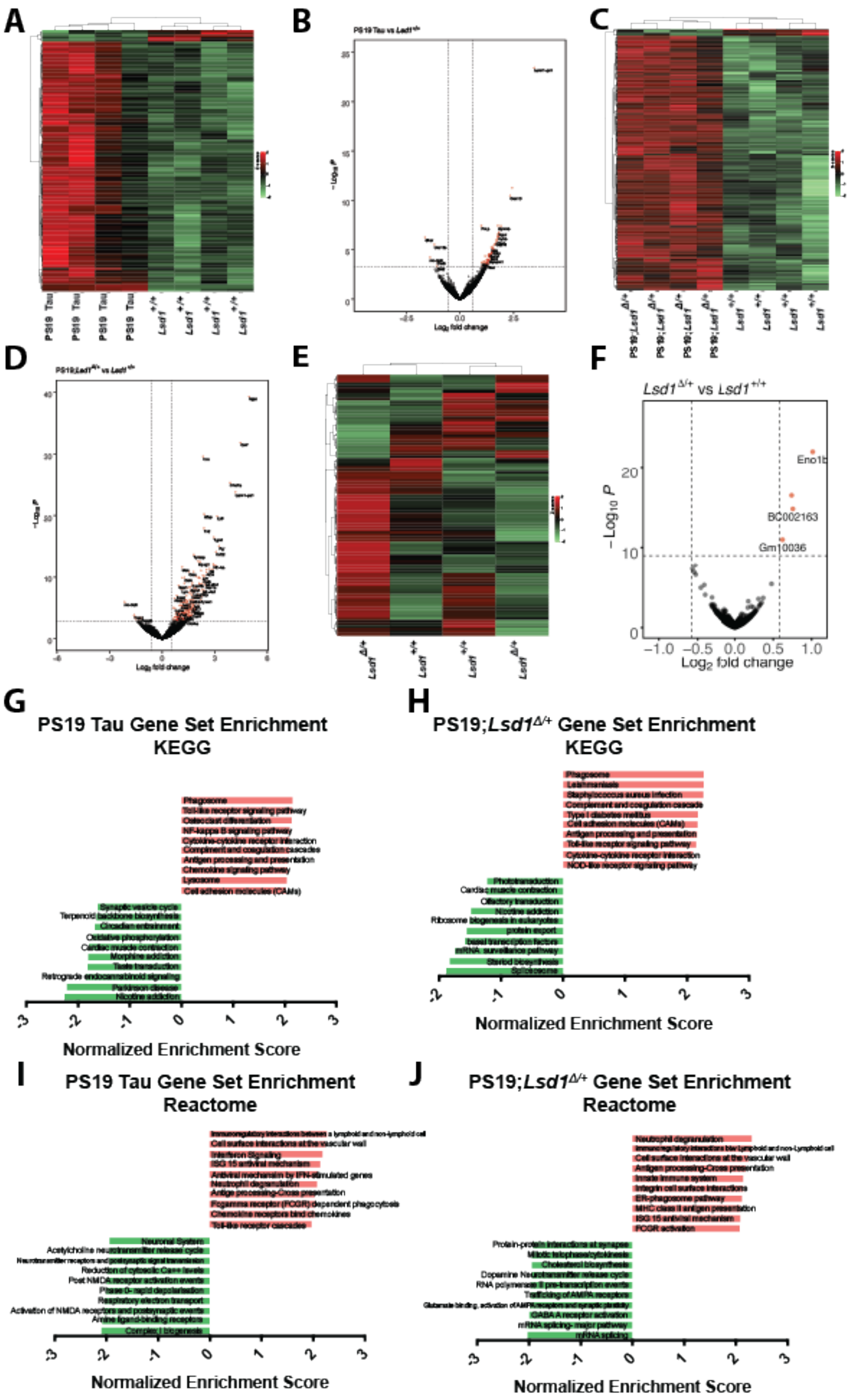

A
B
C
D
E
F
G
H
I
J
PS19 Tau
Lsd1 +/+
PS19;Lsd1 Δ/+
Lsd1 Δ/+
Lsd1 Δ/+ vs Lsd1 +/+
Eno1b
BC002163
Gm10036
−Log10 P
Log2 fold change
PS19 Tau Gene Set Enrichment KEGG
Phagosome
Toll-like receptor signaling pathway
Osteoclast differentiation
NF-kappa B signaling pathway
Cytokine-cytokine receptor interaction
Compliment and coagulation cascades
Antigen processing and presentation
Chemokine signaling pathway
Lysosome
Cell adhesion molecules (CAMs)
Synaptic vesicle cycle
Terpenoid backbone biosynthesis
Circadian entrainment
Oxidative phosphorylation
Cardiac muscle contraction
Morphine addiction
Taste transduction
Retrograde endocannabinoid signaling
Parkinson disease
Nicotine addiction
Normalized Enrichment Score
PS19;Lsd1Δ/+ Gene Set Enrichment KEGG
Phagosome
Leishmaniasis
Staphylococcus aureus infection
Complement and coagulation cascade
Type I diabetes mellitus
Cell adhesion molecules (CAMs)
Antigen processing and presentation
Toll-like receptor signaling pathway
Cytokine-cytokine receptor interaction
NOD-like receptor signaling pathway
Phototransduction
Cardiac muscle contraction
Olfactory transduction
Nicotine addiction
Ribosome biogenesis in eukaryotes
protein export
basal transcription factors
mRNA surveillance pathway
Steroid biosynthesis
Spliceosome
Normalized Enrichment Score
PS19 Tau Gene Set Enrichment Reactome
Immunoregulatory interactions between a lymphoid and non-lymphoid cell
Cell surface interactions at the vascular wall
Interferon Signaling
ISG 15 antiviral mechanism
Antiviral mechanism by IFN-stimulated genes
Neutrophil degranulation
Antigen processing-Cross presentation
Fcgamma receptor (FCGR) dependent phagocytosis
Chemokine receptors bind chemokines
Toll-like receptor cascades
Neuronal System
Acetylcholine neurotransmitter release cycle
Neurotransmitter receptors and postsynaptic signal transmission
Reduction of cytosolic Ca++ levels
Post NMDA receptor activation events
Phase 0- rapid depolarisation
Respiratory electron transport
Activation of NMDA receptors and postsynaptic events
Amine ligand-binding receptors
Complex I biogenesis
Normalized Enrichment Score
PS19;Lsd1Δ/+ Gene Set Enrichment Reactome
Neutrophil degranulation
Immunoregulatory interactions b/w Lymphoid and non-Lymphoid cell
Cell surface interactions at the vascular wall
Antigen processing-Cross presentation
Innate Immune system
Integrin cell surface interactions
ER-phagosome pathway
MHC class II antigen presentation
ISG 15 antiviral mechanism
FCGR activation
Protein-protein interactions at synapse
Mitotic telophase/cytokinesis
Cholesterol biosynthesis
Dopamine Neurotransmitter release cycle
RNA polymerase II pre-transcription events
Trafficking of AMPA receptors
Glutamate binding, activation of AMPA receptors and synaptic plasticity
GABA A receptor activation
mRNA splicing- major pathway
mRNA splicing
Normalized Enrichment Score

**Fig. S8: Differential expression in 9 month old *Lsd1*$^{\Delta/+}$, PS19 Tau, and PS19;*Lsd1*$^{\Delta/+}$ hippocampus. A,C,E,** Heatmap of differentially expressed RNA-seq transcripts between *Lsd1*$^{+/+}$ (*n*=4) and PS19 Tau (*n*=4) (**A**), PS19; *Lsd1*$^{\Delta/+}$ (*n*=4) (**C**), and *Lsd1*$^{\Delta/+}$ (*n*=2) (**E**) mouse hippocampus. Samples are hierarchically clustered by relative expression of differentially expressed transcripts. Relative higher (red) and lower (green) expression is indicated. **B,D,F,** Volcano plot of $\log_2$ fold-changes in gene expression (x-axis) by statistical significance (-$\text{Log}_{10}$ P-value; y-axis) in PS19 Tau (**B**), PS19; *Lsd1*$^{\Delta/+}$ (**D**), and *Lsd1*$^{\Delta/+}$ (**F**) compared to *Lsd1*$^{+/+}$ mouse hippocampus. Each dot represents a transcript, and the dotted line represents a significance $\log_2$ fold change cut off of 0.5. **G-J,** Histogram of Gene Set Enrichment Analysis compared to KEGG pathways (**G,H**) and the Reactome (**I,J**). The top ten most enriched (red) and depleted (green) gene sets in the PS19 Tau (**G,I**) and PS19; *Lsd1*$^{\Delta/+}$ (**H,J**) are shown with normalized enrichment scores.
