## supplemental figures 9-11 for "The inhibition of LSD1 via sequestration contributes to tau-mediated neurodegeneration"

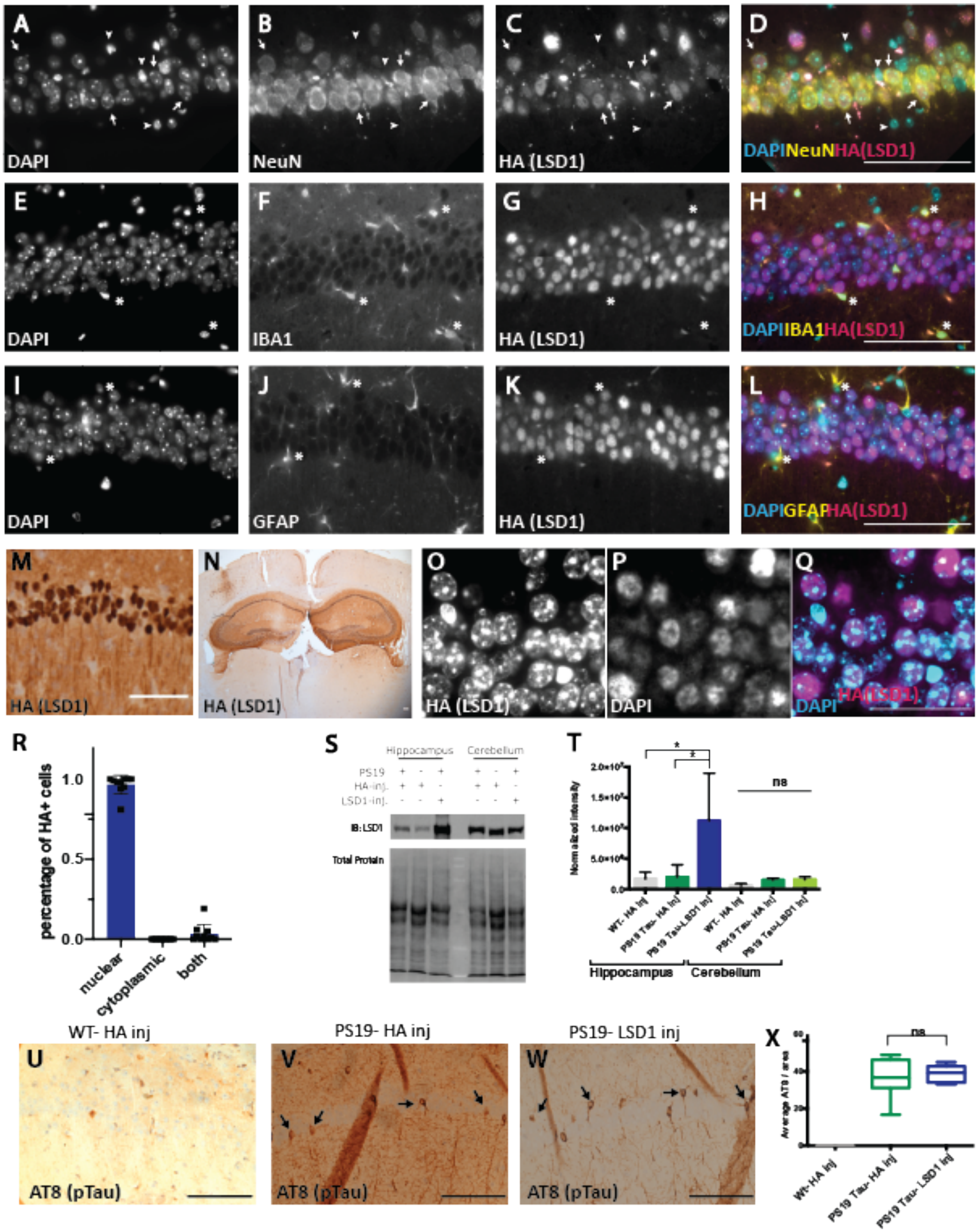
A
B
C
D
DAPI
NeuN
HA (LSD1)
DAPINeuNHA(LSD1)
E
F
G
H
DAPI
IBA1
HA (LSD1)
DAPIIBA1HA(LSD1)
I
J
K
L
DAPI
GFAP
HA (LSD1)
DAPIGFAPHA(LSD1)
M
N
O
P
Q
HA (LSD1)
HA (LSD1)
HA (LSD1)
DAPI
DAPI HA(LSD1)
R
percentage of HA+ cells
1.0
0.5
0.0
nuclear
cytoplasmic
both
S
Hippocampus
Cerebellum
PS19
HA-inj.
LSD1-inj.
IB: LSD1
Total Protein
T
Normalized intensity
ns
WT- HA inj
PS19 Tau- HA inj
PS19 Tau-LSD1 inj
WT- HA inj
PS19 Tau- HA inj
PS19 Tau-LSD1 inj
Hippocampus
Cerebellum
WT- HA inj
PS19- HA inj
PS19- LSD1 inj
U
V
W
AT8 (pTau)
AT8 (pTau)
AT8 (pTau)
X
ns
Average AT8 / area
60
40
20
0
Wt- HA inj
PS19 Tau- HA inj
PS19 Tau- LSD1 inj

**Fig. S9: LSD1 overexpression in hippocampal neurons of PS19 Tau mice. A-D,** Representative immunofluorescence labeling in a WT- HA inj mouse showing DAPI (**A**), NeuN (**b**B, HA (which represents the LSD1 virus, hereafter denoted as HA(LSD1)) (**C**), and merged (**D**). Viral produced LSD1 is present in NeuN+ neurons. Arrows denote NeuN+ cells that have HA expression. Arrowheads denote cells that lack NeuN staining and also lack HA expression. **E-H,** Representative immunofluorescence labeling showing DAPI (**E**), IBA1 (**F**), HA(LSD1) (**G**), and merged (**H**). Asterisks denote cells stained positive for IBA1 (**E-H**), which lack HA expression. **I-L,** Representative immunofluorescence labeling showing DAPI (**I**), GFAP (**J**), HA(LSD1) (**K**), and merged (**L**) images. Asterisks denote cells stained positive for GFAP (**I-L**), which lack HA expression. **M-N,** Immunohistochemistry staining for HA(LSD1) showing expression localized to the nucleus of neurons (**M**) specifically within the hippocampus (**N**). Scale bars=50µm. **O-Q,** Representative immunofluorescence labeling of HA tagged LSD1 in 11 month old PS19-LSD1 inj mouse. DAPI (**O**), HA (LSD1) (**P**), and merged (**Q**) showing that viral produced LSD1 continues to be expressed and localized to the nucleus at the time of rescue, when LSD1 is normally becoming sequestered to the cytoplasm. Scale bars=25µm. **R,** quantification of HA(LSD1) localization shown in **O-Q.** HA (LSD1) was scored as being localized to the nucleus, the cytoplasm, or both areas. Values are mean ± SD, (*n*=10) one-way analysis of variance (ANOVA) with Tukey's post hoc test *P<0.05, **P<0.01,****P<0.001. **S,** Representative image of immunoblot for LSD1 protein and corresponding total protein blot in the hippocampus versus the cortex of mice injected with either LSD1 or HA only expressing virus. **T,** Quantification of immunoblot for LSD1 normalized to total protein loaded per sample shows overexpression in the hippocampus, but not the cortex. Values are mean ± SD (*n*=3, one-way analysis of variance (ANOVA) with Tukey's post hoc test (two-sided), *P<0.05, ns=not significant. **U-W,** Representative image of immunohistochemistry staining of phosphorylated tau (AT8 antibody) in the CA1 region of the hippocampus in 11 month old WT- HA inj (**U**), PS19- HA inj **(V),** and PS19- LSD1inj (**W**) mice. Arrows denote AT8 positive immunoreactivity. Scale bars=50µm. (**X**) Quantification of average AT8 positive tau immunoreactivity per area from histology represented in **U-W**. Box plot edges are 25$^{th}$ and 75$^{th}$ percentile, central line is the median, and whiskers are max and min (*n*=8, one-way analysis of variance (ANOVA) with Tukey's post hoc test (two-sided), ns=not significant).

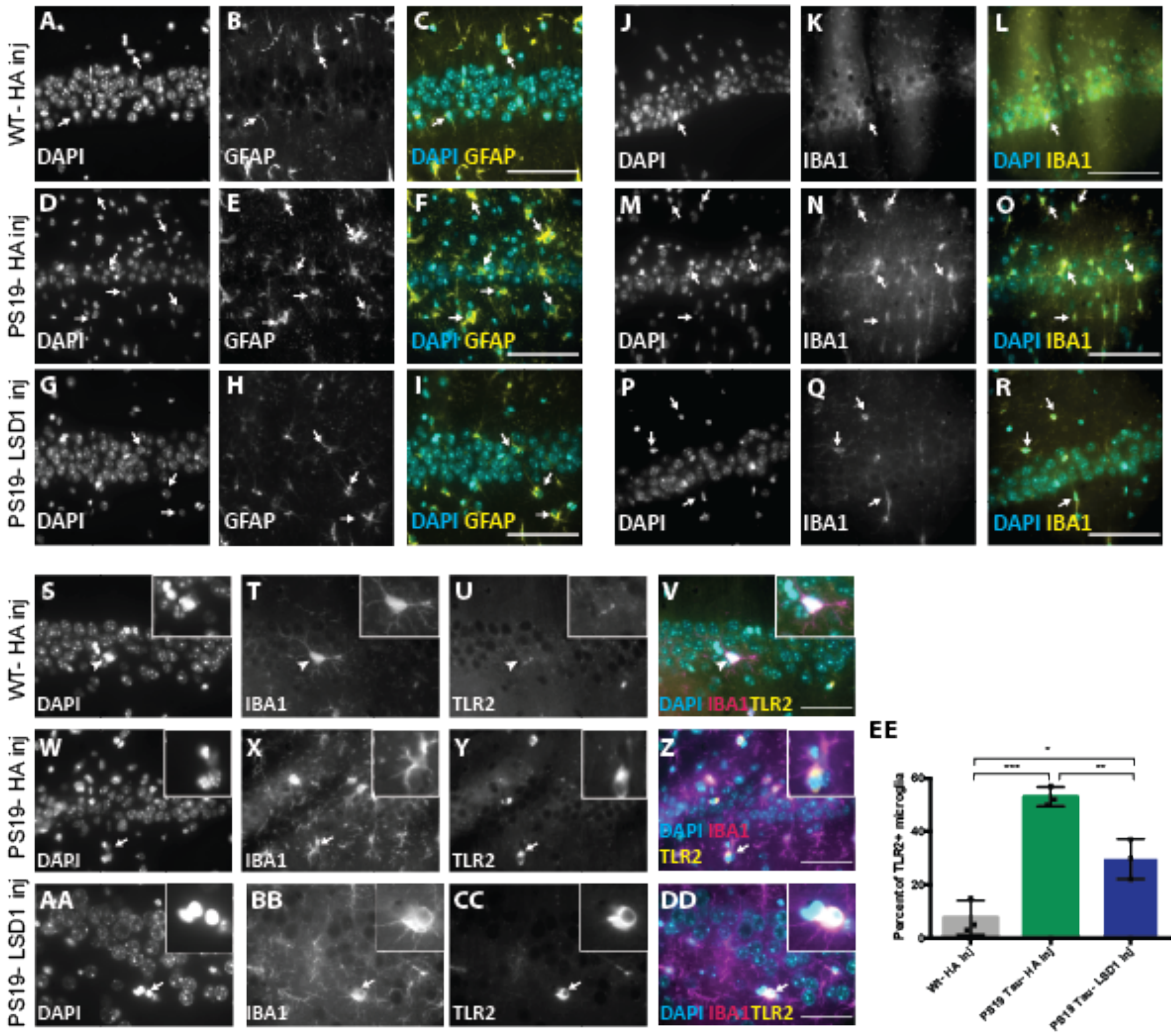


**Fig. S10: LSD1 overexpression reduces the gliosis in PS19 Tau mice. A-I,** Representative immunofluorescence showing DAPI (**A,D,G**), astrocyte marker GFAP (**B,E,H**), and merged (**C,D,I**) images in WT- HA inj (**A-C**), PS19-HA inj (**D-F**), and PS19- LSD1 inj (**G-I**). Arrows denote GFAP+ astrocytes. **J-R,** Representative immunofluorescence showing DAPI (**J,M,P**), microglia marker IBA1 (**K,N,Q**), and merged (**L,O,R**) images in WT-HA inj (**J-L**), PS19-HA inj (**M-N**), and PS19- LSD1 inj (**P-R**). Arrows denote IBA1+ microglia. **S-DD,** Representative immunofluorescence labeling showing DAPI (**S,W,AA**), microglia marker IBA1 (**T,X,BB**), activated microglia marker TRL2 (**U,Y,CC**), and merged (**Y,Z,DD**) images in WT-Ha inj (**S-V**), PS19-HA inj (**W-Z**), and PS19-LSD1 inj (**AA-DD)**. Inset of microglia that is IBA1 positive but TRL2 negative (**S-V,** denoted by arrowhead) or both IBA1 and TRL2 positive (**W-DD**, denoted by arrow). All images are from the CA1 region of the hippocampus Scale bars=50µm. **EE,** Quantification of the percentage of microglia that are TRL2+ in WT- HA inj, PS19- HA inj, and PS19- LSD1 inj mice represented in **S-DD**. Values are mean ± SD are (*n*=3, one- way analysis of variance (ANOVA) with Tukey's post hoc test, *p<0.05, **p<0.01, ***p<0.005).

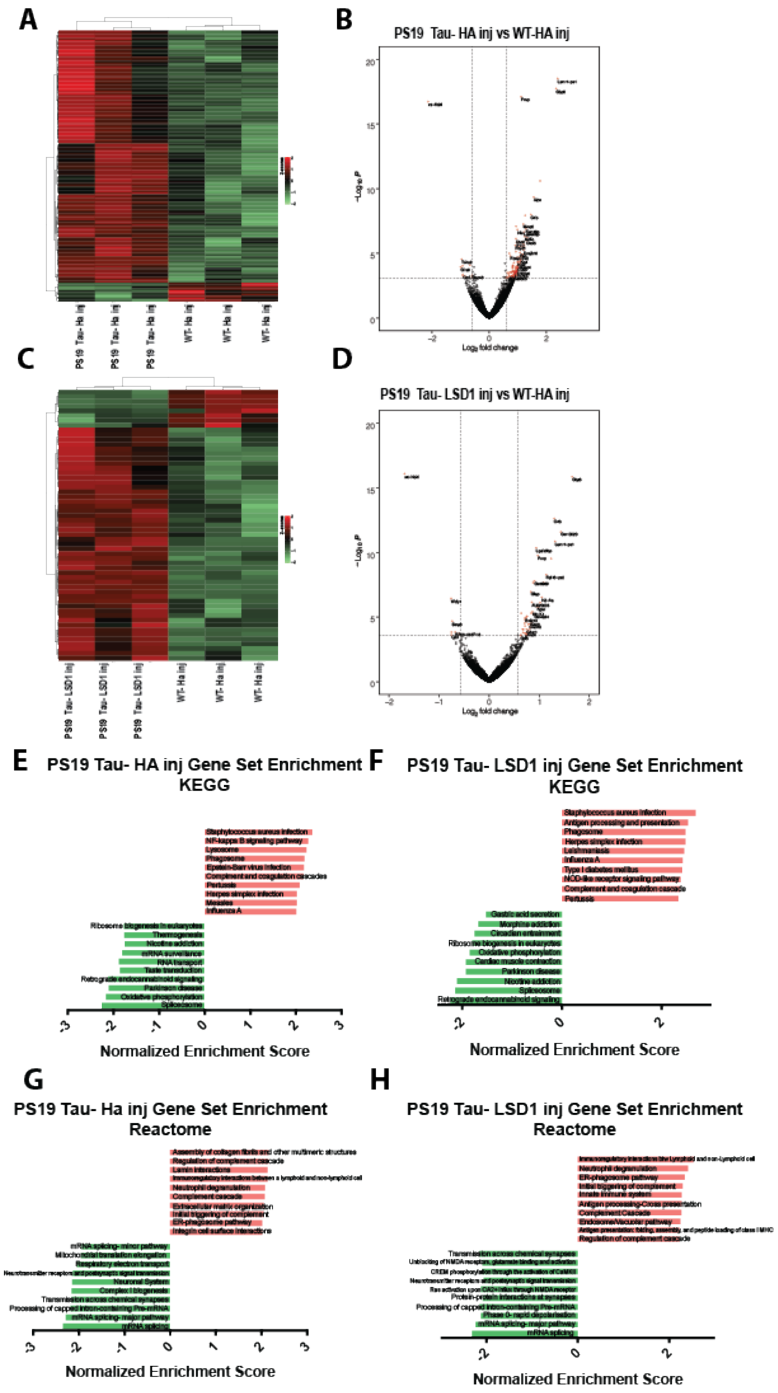


**Fig. S11: Differential expression in 11 month viral injected PS19 Tau mice. A,C,**, Heatmap of differentially expressed RNA-seq transcripts in PS19 Tau- HA inj (**A**) and PS19 Tau- LSD1 inj (**C**) versus WT-HA inj mouse hippocampus. Samples are hierarchically clustered by relative expression of differentially expressed transcripts. Relative higher (red) and lower (green) expression is indicated. **B,D,** Volcano plot of $\log_2$ fold-changes in gene expression (x-axis) by statistical significance ($-\text{Log}_{10}$ P-value; y-axis) in PS19 Tau- HA inj (**B**) and PS19 Tau- LSD1 inj (**D**) compared to WT- HA inj mouse hippocampus. Each dot represents a transcript, and the dotted line represents a significance $\log_2$ fold change cut off of 0.5. **E-H,** Histogram of Gene Set Enrichment Analysis compared to KEGG pathways (**E,F**) and the Reactome (**G,H**)**.** The top ten most enriched (red) and depleted (green) gene sets in the PS19 Tau- HA inj (**E,G**) and PS19 Tau- LSD1 inj (**F,H)** are shown with normalized enrichment scores.
