## supplemental table 1 and movie 1-4 figure legends for "The inhibition of LSD1 via sequestration contributes to tau-mediated neurodegeneration"

| Target | Manufacturer | Clone | Lot number | Experiment | Dilution |
|---|---|---|---|---|---|
| NeuN | Millipore MAB377 | A60 | 2392283 | Mouse IF | 1:500 |
| LSD1 | Abcam 17721 | | GR3193508-2 | Mouse IF | 1:100 |
| PHF-Tau | ThermoFisher MN 1020 | AT8 | TI2611431 | Mouse IHC | 1:1,000 |
| | | AT8 | TI2611431 | Mouse IF | 1:200 |
| HA | Abcam ab130275 | 16B12 | GR3190856-12 | Mouse IHC | 1:500 |
| | | 16B12 | GR3190856-12 | Mouse IF | 1:100 |
| HA | Abcam ab9110 | | GF3224022-1 | Mouse IF | 1:500 |
| GFAP | Dako Z0334 | | 20047046 | Mouse IHC | 1:100 |
| IBA1 | Synaptic System 234004 | | 2-16 | Mouse IF | 1:100 |
| Neurofilament (phospho) | Millipore NE1022 | SMI-31R | | Mouse IHC | 1:500 |
| PHF-1 | Peter Davies (Albert Einstein College of Medicine, New York, NY) | | | Mouse IHC | 1:1,000 |
| LSD1 | Cell Signaling 2139 | | 5 | Immunoblot | 1:1,000 |
| TLR2 | Abcam ab9100 | TL2.1 | GR3189369-7 | Mouse IF | 1:100 |

**Table S1. Primary antibodies used for immunohistochemistry (IHC) and immunofluorescence (IF) experiments.** Shown for each antibody are the target antigen, manufacturer, experiments used and corresponding experimental dilution.

**Movie S1. Reduction of LSD1 in PS19 Tau mice exacerbates paralysis. (0-0:06 sec)**, Tail suspension of 12 month old *Lsd1*$^{\Delta/+}$ mouse that has full mobility in hindlimbs and no paralysis. **(0:07-0:13 sec)**, Tail suspension of 12 month old PS19 Tau mouse that has a hindlimb clasp, seen by holding legs contracted inward and not moving them, but is still mobile upon release. **(0:14-0:19 sec)**, Tail suspension of 12 month old PS19;*Lsd1*$^{\Delta/+}$ mouse that is terminally paralyzed.

**Movie S2. Magnetic Resonance Imaging of hippocampal atrophy throughout the brain when LSD1 is reduced in PS19 Tau mice. (0-0:33),** Serial coronal slices from T2-weighted RARE MRI, starting posterior ending anteriorly, through the brain of a 6 month old *Lsd1*$^{\Delta/+}$ mouse (**0:03-0:13**), PS19 Tau mouse (**0:13-0:22**), and PS19;*Lsd1*$^{\Delta/+}$ mouse (**0:23-0:32**). (**0:33-1:05**), Serial coronal slices from T2-weighted RARE MRI, starting posterior ending anteriorly, through the brain of the same mouse *Lsd1*$^{\Delta/+}$ mouse (**0:35-0:45**), PS19 Tau mouse (**0:45-0:55**), and PS19;*Lsd1*$^{\Delta/+}$ mouse (**0:55-1:05**) at 10 months old. High intensity areas (white/light grey) signify ventricular dilatation.

**Movie S3. Hippocampal injection of viral LSD1 did not affect development of paralysis in PS19 mice. (0-0:13 sec)**, Tail suspension of 11 month old WT- HA inj mouse that has full mobility in hindlimbs and no paralysis. **(0:14-0:28 sec)**, Tail suspension of 11 month old PS19- HA mouse that has a hindlimb clasp, seen by holding legs contracted inward and not moving them, but is still mobile upon release. **(0:29-37sec)**, Tail suspension of 11 month old HA- LSD1 inj mouse that similarly has a hindlimb clasp holding legs contracted inward and not moving them, but is still mobile upon release.

**Movie S4. Pathological Tau Induces Neurodegeneration by sequestering and inhibiting LSD1** In healthy hippocampal and cortical neurons, LSD1 is translated in the cytoplasm and transported through the nuclear pore into the nucleus where it is continuously required to repress inappropriate transcription. In tauopathy, LSD1 is translated in the cytoplasm, but as pathological tau accumulates in the cytoplasm it blocks LSD1 from being imported into the nucleus. This interferes with the continuous requirement for LSD1, resulting in neuronal cell death.

**Data S1. (separate file)**

**Expression changes in 9 month old *Lsd1*$^{\Delta/+}$, PS19 Tau, PS19;*Lsd1*$^{\Delta/+}$ mice.** Spreadsheets for all, significantly upregulated, and significantly downregulated transcripts. Provided for each genotype with two biological replicates are the log2 fold change, P-value, and p adjusted-value as determined by DESEQ2.
