## Supplementary material for "The inhibition of LSD1 via sequestration contributes to tau-mediated neurodegeneration": Data S2

### Expanded Methods for RNA Sequencing Analysis

#### Quality Control

Sequencing data (FASTQ files) were uploaded to the Galaxy web platform, (*public server at http://*usegalaxy.org). Paired-end data from each sample (forward and reverse) were put together into a data collection (2 files per collection). *Quality assessment on each data collection of FASTQ files was performed using* FASTQC (v.0.11.7). FASTQC returned many quality assessment metrics. The sequence quality histogram illustrated the mean quality (or Phread) score per base across the 50bp reads. An average Phread score of 20 or greater and minimum length of 36 base pairs after trimming was required to retain the read for further downstream analysis. FASTQC also provided information such as sequence length distribution, sequence duplication levels, overrepresented sequences and adapter content. We ensured these metrics were within the acceptable range and no adapter content was present. Sequence trimming was performed using Trimmomatic (v.0.36.5). Default settings were used unless noted otherwise below. Trimmomatic parameters utilized within the Galaxy web platform:

- select “paired-end” (as collection)
- “Perform initial ILLUMINACLIP step?” = NO
- Use default setting for 1st trimmomatic operation
- Sliding window trimming, Number of bases to average across = 4,
- Average quality score required = 20
- Insert 2nd trimmomatic operation
- from dropdown list select “minlen”
- minimum length of reads to be kept: 36bp

#### Sequence Mapping

Paired-end reads were mapped to the GRCm38 genome using HISAT2 (v.2.1.0). GRCm38 (DNA primary assembly) FASTA file was downloaded from the Ensembl FTP site on 7/24/18 and uploaded to the galaxy web platform. Default settings were used unless noted otherwise below. HISAT2 parameters utilized within the Galaxy web platform:

- Select ‘use genome from history”
- Select uploaded GRCm38 DNA primary assembly FASTA file
- “Paired-end reads”
- “Unstranded”
- Under ‘summary options’ select ‘output summary’ and ‘print summary to file’

**Sequence Mapping Quality Control**

Unmapped, unpaired and multiply mapped reads were then removed using Filter SAM or BAM (v.1.1.2). Filter SAM or BAM parameters utilized within the Galaxy web platform:

- Select BAM file generated by HISAT2
- Set “minimum MAPQ quality score” to 40
- Set “filter on bitwise flag” to yes
- Select “read is paired” and “read is mapped in a proper pair”

HISAT2 MAPQ scores range from 0-60:
60 - uniquely mapped read, regardless of number of mismatches / indels
1 - multiply mapped, perfect match or few mismatches / indels
0 - unmapped, or multiply mapped and with lots of mismatches / indels

Utilizing a minimum MAPQ quality score > 40 allowed for the removal of non-uniquely mapped reads. Additionally, BAM files from the Filter SAM or BAM utility were analyzed for mapping quality using Flagstat.

**Differential Expression**

Assignment of transcripts to GRCm38 genomic features was performed using Featurecounts (v.1.6.0.6). GRCm38.93 GTF file was downloaded from Ensembl FTP site on 7/21/18 and uploaded to the galaxy web platform. Featurecounts parameters utilized within the Galaxy web platform:

- Select BAM files generated by Filter SAM or BAM utility
- "Unstranded"
- Select annotation file "in your history"
- Select Ensembl GTF file previously uploaded
- "Output format": DESEQ2 compatible
- Select yes for "create gene length file" (needed calculate TPM or FPKM)
- Under "options for paired-end reads" menu, enable 'count fragments instead of reads"

Differentially expressed transcripts were determined using DESEQ2 (v.2.11.40.2). Tabular count files from Featurecounts were used to analyze differential expression between genotypes. Within DESEQ2, the factor was genotype with the different levels being experimental genotype vs control/ WT genotype. The analysis was run independently for each experimental-control pair. DESEQ2 was also utilized to generate normalized count tables. These counts represent expression values normalized for differences in sequencing depth and composition bias to allow for direct comparison between samples via heatmap visualizations. DESEQ2 parameters utilized within the Galaxy web platform:

- Select output from Featurecounts utility (.tabular count files)
- Factor = genotype
- Factor level 1 = experimental samples of the same genotype
- Factor level 2 = control samples
- Repeat for each genotype compared to control
- Under "Files have header?" select "yes" (output from featurecounts have a header)
- Under "Output normalized counts table?" select "yes"

#### RNA Sequencing Data Processing

##### Transcripts per million reads (TPM) expression values

The following R script was used to determine gene expression in terms of transcripts per million reads (TPM). Raw count files and the feature length file from Featurecounts was the input data. The analysis was performed independently for each genotype. The following is a representative example.

```
#TPM = (read count / gene length in kilobases) / (sum of count/gene length column / 10^6)
## Read in Lsd1 control raw counts files and feature length file
controla.counts <- tbl_df(read.delim("controlA_SRR5535973.featurecounts.tabular",
                              stringsAsFactors = FALSE, header = TRUE))
controlb.counts <- tbl_df(read.delim("controlB_SRR5535974.featurecounts.tabular",
                              stringsAsFactors = FALSE, header = TRUE))
control.lengths <- tbl_df(read.delim("controlB_SRR5535974.featurelengths.tabular",
                               stringsAsFactors = FALSE, header = TRUE))

#Combine dataframes by Ensembl IDs
control <- control.lengths %>%
  left_join(controla.counts,by = "Geneid" ) %>%
  left_join(controlb.counts,by = "Geneid") %>%

#Convert feature lengths column from base pairs to kb
  mutate(control.length.kb = Length/1000) %>%
  rename(controlb.count = SRR5535974_controlB_.fastqsanger,
       controla.count = SRR5535973_controlA_.fastqsanger) %>%

#Calculate RPK for each sample
  mutate(controlaRPK = controla.count/control.length.kb,
       controlbRPK = controlb.count/control.length.kb)

#Calculate scaling factor for each sample
controla.scalingfactor <- sum(control$controlaRPK)/1000000
controlb.scalingfactor <- sum(control$controlbRPK)/1000000

#Calculate TPM for each sample
control <- control %>%
  mutate(controlaTPM = controlaRPK/controla.scalingfactor,
             controlbTPM = controlbRPK/controlb.scalingfactor) %>%

#Calculate mean value for both samples
  mutate(avg.controlTPM = rowMeans(select(.,controlaTPM,controlbTPM))) %>%
  arrange(desc(avg.controlTPM))
```

**Identification of differentially expressed genes (DEGs)**

For all datasets, a cutoff of adjusted p-value < 0.3 and abs ($log_2$ fold change) > 0.58 was applied.

This was performed using the following R script.

```
#Load Tidyverse package
library(tidyverse)
```

```
#Read in DESEQ2 results file
sample <- tbl_df(read.delim("Galaxy1-
        [DESeq2_result_file_on_LSD1mutant_vs_LSD1control].tabular", header = FALSE, sep
        = "\t", stringsAsFactors = FALSE))

#Add column identifiers
colnames(sample) <- c("ensembl.geneid","base.mean","log2fc","stderr","wald-
        stats","p.value","p.adj")

#Filter out genes with NA values for P.adjusted or log2 Fold-Change
#Apply p-adjusted and Log2 Fold-Change cutoffs
#Arrange in descending order by log2fc values
DEGs <- sample %>%
        select(ensembl.geneid, log2fc, p.adj, p.value ) %>%
        filter(!is.na(p.adj) & !is.na(log2fc)) %>%
        filter(p.adj < 0.3 & abs(log2fc) > 0.58 ) %>%
        arrange(desc(log2fc))

#Output CSV file with list of DEGs and corresponding gene symbols
write.csv(DEGs,"lsd1mut.DEGs.csv", row.names = FALSE)
```

**Convert Ensembl IDs to Gene Symbols**

The following R script was used to match mouse Ensembl Gene IDs in expression data to corresponding mouse gene symbols.

```
## install GenomicFeatures and EnsDb.Mmusculus.v79 package to read in mouse gene
## identifiers
BiocManager::install ("GenomicFeatures")
BiocManager::install ("EnsDb.Mmusculus.v79")
library(GenomicFeatures)
library (EnsDb.Mmusculus.v79)

## Pull gene identification data from Ensembl Database
keys <- keys(EnsDb.Mmusculus.v79)
anno.result <- select(EnsDb.Mmusculus.v79, keys=keys,
        columns=c("GENEID","SYMBOL","GENENAME","ENTREZID"),keytype="GENEID)

## Subset annotation file for Ensembl ID and Gene Symbol
annot <- select(anno.result, GENEID, SYMBOL)

## Join annotation table and sample gene expression data by Ensembl IDs
Sample.symbol <- left_join(sample, annot, by = c("ensembl.geneid" = "GENEID"))
```

**Heatmaps**

Heatmaps and hierarchical clustering were generated using the “complex heatmap package” in R. Input was normalized counts files from DESEQ2. Example R Script:

```
#read in deseq2 results file
sample1.counts <- read.delim("Galaxy204-[Normalized_counts_file_on_.tabular",
                              header = TRUE, sep = "\t", stringsAsFactors = FALSE)

#assign column header names
colnames(sample1.counts)  <- c("geneid", "sample1", "sample2","sample3", "sample4" )

#read in csv with DEGs
sample1.DEGs <- tbl_df(read.csv("sample1.DEGs.csv", stringsAsFactors = FALSE))

#Create matrix for generating heatmap
samples <- sample1.counts %>%

#join dataframes by geneid
#Subset normalized counts file for DEGs
        inner_join(sample1.DEGs, by = "geneid") %>%
        arrange(p.value) %>%

#log2 transform count data
        mutate("sample.1"= log2(sample1 + 1), "sample.2"= log2(sample2 +1),
        "sample.3"= log2(sample3 +1), "sample.4"= log2(sample4 +1)) %>%

#subset data to only contain the log2 transformed counts
        select("sample.1", "sample.2", "sample.3", "sample.4") %>%

# convert dataframe to matrix
        data.matrix()

#Install the “Complex Heatmap” package and “circlize” package (for colors)
install_github("jokergoo/ComplexHeatmap")
install.packages("circlize")
library(ComplexHeatmap)
library(circlize)

#Set heatmap colors
col_fun = colorRamp2(c(-2, 0, 2), c("light green", "black", "red"))

#Scale data
```

```
sample_scaled = t(scale(t(samples)))

#Generate final heatmap
#Cluster both rows and columns
#Hierarchical Clustering method = Complete
#Hierarchical Clustering distance algorithm = pairwise distance function
Heatmap(samples,
        name = "samples",
        col = col_fun,
        cluster_rows = TRUE,
        cluster_columns = TRUE,
        clustering_method_rows = "complete",
        clustering_method_columns = "complete",
        clustering_distance_rows = function(x, y) 1 - cor(x, y),
        clustering_distance_columns = function(x, y) 1 - cor(x, y),
        heatmap_legend_param = list (title = 'Z-scores',
                   legend_height = unit(4, "cm"),
                  title_position = "lefttop-rot"
    ))
```

**Volcano plots**

Volcano plots were produced using the "Enhanced Volcano" package in R. A volcano plot is a scatterplot for the visual comparison of statistical significance (-log10 P-value, y axis) and magnitude of change (Log2 Fold-Change values, x axis). Example R Script:

```
# Install and load Enhanced Volcano package
BiocManager::install("EnhancedVolcano", version = "devel")
library(EnhancedVolcano)

## Enhanced volcano only seems to work when you read in a tab delim text file (not read.csv()
## or tbl_df(read.csv))
## Read in deseq2 output file
lsd1mut2 <- read.delim("lsd1mut.gs.rmna.txt", header = TRUE, sep = "\t", dec = ".")

# Plot volcano plot
plot2 <- EnhancedVolcano( lsd1mut2,
                  lab = lsd1mut2$gene.symbol,
                  x = "log2fc",
                  y = "p.value",
                  pCutoff = 0.007131771,
                  FCcutoff = 2,
                  xlab = bquote(~Log[2]~ "fold change"),
                  ylab = bquote(~-Log[10]~italic(P)),
                  transcriptPointSize = 1.5,
```

```
transcriptLabSize = 3.0,
title = "LSD1 cKO vs WT",
col = c("black", "black", "black", "red3"),
colAlpha = 0.5,
xlim = c(-8,8.5),
ylim = c(0,65),
cutoffLineType = "dashed",
cutoffLineCol = "black",
cutoffLineWidth = 0.5,
legendPosition = "none",
DrawConnectors = FALSE,
widthConnectors = 0.2,
colConnectors = "grey30",
border = "full",
borderWidth = 1.5,
borderColour = "black",
gridlines.major = FALSE,
gridlines.minor = FALSE)
```

**Gene Set Enrichment Analysis (GSEA)**

Gene Set Enrichment Analysis (Pre-ranked list) was performed using the online platform WebGestalt. The metric used for determining rank was the standard: Sign ($log_2$ Fold-Change) * -$log_{10}$ (P-Value). The ranked list file (*.rnk) was created using the following r script.

```
## Load tidyverse package
library(tidyverse)

## Read in NCBI Homologene data
hom <- read.table( "homologene.data", sep = "\t", header = TRUE, quote = "", stringsAsFactors
= FALSE)

## Read PS19_Lsd1 deseq2 output
ps19_lsd1 <- read.csv("transhet.gs.rmna.csv", stringsAsFactors = FALSE)

## subset homologene table
#mouseId = 10090
#humanId = 9606
```

```
## subset homologene table for taxid and human gene symbol
homology.human <- hom %>%
  filter(taxId == "9606") %>%
  select(id, geneSymbol) %>%
  rename(geneid_human = geneSymbol)

## subset homologene table for taxid and mouse gene symbol
 homology.mouse <- hom %>%
  filter(taxId == "10090" ) %>%
  select(id, geneSymbol) %>%
  rename(geneid_mouse = geneSymbol)

## Add column of taxids corresponding to mouse gene symbols
x <- ps19_lsd1 %>%
  left_join(homology.mouse, by = c("gene.symbol" = "geneid_mouse")) %>%

## Add column of human gene symbols corresponding to taxids
  left_join(homology.human, by = "id") %>%

## Include only log2fc, p value and human gene symbols columns and filter out NA values
  select(log2fc, p.value, geneid_human) %>%
  filter(!is.na(geneid_human), !is.na(log2fc), !is.na(p.value)) %>%

## Calculate metric for GSEA pre-ranked list
  mutate(fcSign = sign(log2fc),
         logP = -log10(p.value),
         metric = logP*fcSign) %>%

## Select human gene symbol and metric (sorted by metric value)
  select(geneid_human, metric) %>%
  arrange(desc(metric))

## Write ranked expression file (.rnk) for running GSEA on Web Gestalt platform
write.table(x, file = "ps19_lsd1.expression.rnk", quote = FALSE, sep = "\t", row.names =
FALSE,  col.names = FALSE)
```

### Data Visualization

#### Scatterplot

R script for creating scatterplot to compare expression changes ($log_2$ Fold-Change) genome-wide between P19 Tau mouse and the PS19;*Lsd1*$^{\Delta/+}$ mouse.

```
library(tidyverse)
## Read in expression data from DESEQ2
tau <- read.csv("tau.gs.rmna.csv", stringsAsFactors = FALSE)
transhet <- read.csv("transhet.gs.rmna.csv", stringsAsFactors = FALSE)

## Join together the data from each genotype
corr4 <- inner_join(transhet, tau, by = "ensembl.geneid")

## create new dataframe with upregulated genes (log2fc > .58 & p.adj < 0.3)
up4 <-transhet %>%
  filter(p.adj < 0.3 & log2fc > 0.58 )

## create new dataframe with downregulated genes (log2fc < -.58 & p.adj < 0.3)
down4 <-transhet %>%
  filter(p.adj < 0.3 & log2fc < -0.58 )

## Add new column for identifying DEGs by color
corr4 <- mutate(corr4, condition = ifelse(ensembl.geneid %in% up4$ensembl.geneid, "up",
                                    ifelse(ensembl.geneid %in% down4$ensembl.geneid,
                                    "down","others")))

## Plot scatterplpot using ggplot2
p4 <- ggplot(corr4, aes(x= log2fc.x, y = log2fc.y, colour= condition), na.rm = TRUE) +
  geom_point(size =1, alpha=.75)   +
  theme(legend.position="none",
        panel.grid.major = element_blank(),
        panel.grid.minor = element_blank(),
        axis.line = element_line(colour = "black"),
        panel.background = element_blank(),
        panel.border = element_rect(colour = "grey", fill=NA),
        plot.title = element_text(hjust = 0.5)) +
  scale_color_manual(
    values = c("up" = "dark red",
            "down" = "dark green",
            "others" = "dark grey"))+
  geom_hline(yintercept=0)+
  geom_vline(xintercept=0)+
  xlim(-2.5,2.5) +
  ylim(-2.5,2.5)+
  xlab(expression(paste("PS19;",italic("Lsd1")^Delta,""^ "/+ ","(", log[2], " fold change)"))) +
  ylab(expression(paste("PS19 (", log[2], " fold change)"))) +
  ggtitle(expression(paste("PS19;", italic("Lsd1")^Delta, ""^ "/+", " vs PS19")))
p4 +
geom_abline(intercept = 0, slope = 1, linetype = "dashed")
```
